# A rugged yet easily navigable fitness landscape of antibiotic resistance

**DOI:** 10.1101/2023.02.27.530293

**Authors:** Andrei Papkou, Lucia Garcia-Pastor, José Antonio Escudero, Andreas Wagner

## Abstract

A fitness landscape is a biological analogue of a physical landscape, in which each genotype occupies a location whose elevation corresponds to fitness. Theoretical models predict that rugged fitness landscapes with multiple peaks should impair Darwinian evolution, because natural selection prevents evolving populations from traversing the valleys that lie between peaks. Experimental tests of this prediction are very limited. Here we combine CRISPR-Cas9 genome editing and deep sequencing to map the fitness landscape of more than 260’000 genotypes of the *E. coli folA* gene in an environment harboring the antibiotic trimethoprim. The *folA* gene encodes the key metabolic enzyme dihydrofolate reductase (DHFR), which is also a target of this antibiotic. With 514 mostly low fitness peaks, the DHFR fitness landscape is rugged. Despite this ruggedness, its highest fitness peaks are easily accessible to evolving populations. Fitness-increasing paths to high fitness peaks are abundant, and individual peaks have large basins of attractions. The basins of different peaks overlap, which renders the outcome of adaptive evolution highly contingent on chance events. In sum, ruggedness need not be an obstacle to Darwinian evolution but can reduce its predictability. If true in general, evolutionary biology and other fields of sciences in which landscapes play an important role may have to re-appraise the complexity of optimization problems on realistic landscapes.

---

The fitness landscape is a nearly century-old foundational concept in evolutionary biology^1^. Its influence extends to multiple other disciplines, including ecology^2^, chemistry^3^, computer science^4,5^, the social sciences^6^, and engineering^7^. It is an analogy to a physical landscape in which individual spatial locations correspond to genotypes and the elevation at each location corresponds to the genotype’s fitness. The best-adapted genotypes occupy the highest peak(s) of such a landscape. A population evolving by natural selection explores such a landscape, and natural selection drives such a population uphill to the nearest peak^1,8,9^.

A fitness landscape can be single-peaked or multi-peaked (“rugged”)^10^. Ruggedness can pose a fundamental challenge to evolution’s ability to find a landscape’s highest peaks, because a population evolving under the influence of natural selection can only travel on *accessible* paths through the landscape, i.e., paths in which each mutational step increases fitness^8,11^.

The reason is that natural selection favors high fitness genotypes, and does not allow a population to traverse any low fitness valleys between a local peak of intermediate fitness and nearby higher fitness peaks^8,12^.

Theory predicts that in highly rugged landscapes, most evolving populations will become trapped at local peaks of low fitness^13^. The relationship between landscape ruggedness and peak accessibility has been studied with various theoretical models developed in the 20^th^ century^14^, such as the NK^15^, House-of-Cards^16^, and Rough Mount Fuji models^17^, which may not capture important features of empirical landscapes^18,19^. Because most research on adaptive landscapes remained theoretical until recent decades^8,10^, we still know little about the ruggedness, and even less about the accessibility of high fitness peaks in empirical fitness landscapes.

Early experimental studies to map empirical landscapes measured the fitness of less than 10^3^ genotypes^20–30^. Later studies mapped up to 10^5^ genotypes that were generated by random mutagenesis of a reference (wild type) genotype. The resulting fitness data is dense around the wild type, but sparse everywhere else due to missing information on double, triple, and higher order mutants^31,32^. In other words, such data do not fulfill the important requirement of *combinatorially completeness*, i.e., that the fitness of all allele combinations at the mutagenized loci must be known to permit an exhaustive search of evolutionary paths. Some studies achieved combinatorial completeness by quantifying biochemical properties that may be correlated with fitness, such as the binding of biological molecules in vitro^33–37^. Only the most recent work created simultaneously large and combinatorially complete fitness landscapes in microorganisms such as *S. cerevisiae* and *E. coli*. However, this work focused on other aspects of landscapes, such as the prevalence of higher order epistasis^38,39^, the molecular principles underlying genotype-phenotype relationship^40^, the allosteric effect of mutations^41^, and the evolution of specificity in protein interactions^42,43^.

Here we address the unresolved and fundamental question of the relationship between ruggedness and peak accessibility by mapping a large and combinatorially complete *in vivo* fitness landscape of the *E.coli folA* gene, which encodes the essential metabolic enzyme and antibiotic resistance protein dihydrofolate reductase (DHFR). We find that this landscape is rugged, but its ruggedness does not preclude evolving populations from accessing high fitness peaks.

## Experimental design and reproducible fitness measurements

We performed CRISPR-Cas9^44^ deep mutagenesis to edit the *folA* gene on the bacterial chromosome, randomizing nine nucleotide positions in a part of the gene that is both conserved and implicated in the evolution of antibiotic resistance (Fig. 1A). The result is a combinatorially complete library of almost 4^9^ (262,144) DNA genotypes. They include all possible (64^3^) combinations of codons that encode three successive amino acids of DHFR (wild type sequence: 26A-27D-28L). Missense mutations at these positions provide high resistance to the clinically important antibiotic trimethoprim, which inhibits DHFR. Combinations of these mutations show non-additive (epistatic) interactions^45,46^.

**Fig. 1.**
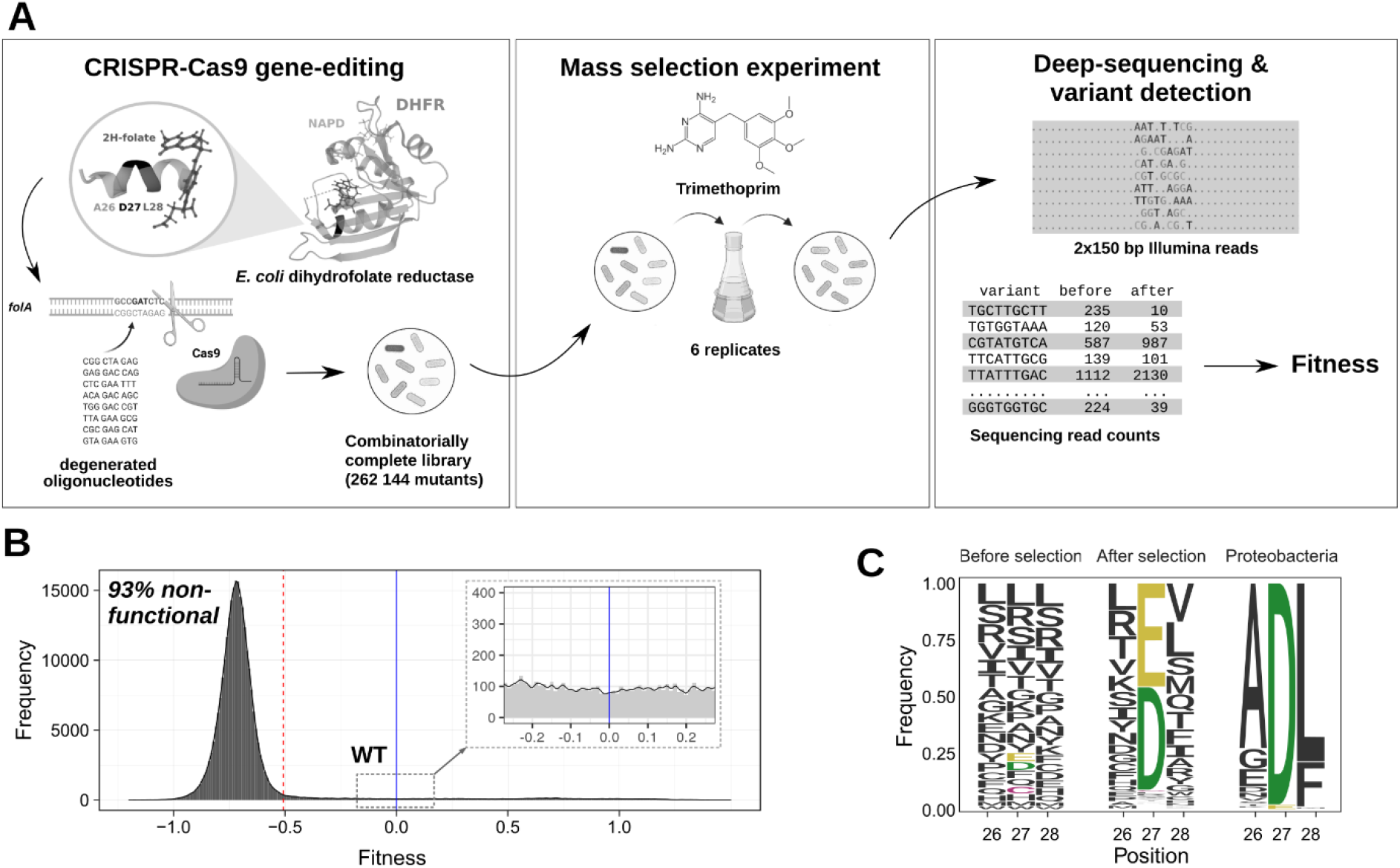
**(A) Experimental design.** (left) We used CRISPR-Cas9 gene editing to create a library of DHFR variants. We targeted a nine bp segment of the *folA* gene on the *E. coli* chromosome, which encodes three amino acids of DHFR (A26-D27-L28). In the folded protein, these three amino acids lie inside the substrate binding pocket. We edited the segment with degenerate oligonucleotides to obtain a library comprising 99.7% of all theoretically possible DHFR genotypes for this segment. (middle) We used this library to perform a mass selection experiment by growing six parallel cultures at the half-inhibitory concentration of trimethoprim (0.4 µg/ml). (right) We amplified the variable DHFR region and performed Illumina paired-end deep sequencing. We determined sequencing read counts for each variant before and after selection and used them to calculate relative fitness (see Methods). **(B) The distribution of fitness effects in the library**. The dashed red line indicates the cut-off value for nonfunctional variants (−0.508). The blue solid line marks the fitness of the wild-type variant, and the insets show the tail of the distribution near the wild type. *N*=261,332 variants. The inset indicates that the distribution has a heavy tail near the wildtype. **(C) The frequency of amino acids at position 26, 27, 28**. The panels show the frequency of amino acids in the library before selection (left panel) and after selection (middle panel). The right panel shows the corresponding amino acid frequencies in DHFRs from proteobacteria, based on an alignment of 5000 orthologous *folA* sequences.

We exposed a population of *E.coli* cells expressing this library to a sublethal dose of trimethoprim, and measured the fitness of library members through deep sequencing in a six-fold replicated mass-selection experiment. The resulting data comprise *folA* variant frequencies before and after selection for 99.7% (261,382/262,144) of all possible variants (Methods). Variant frequencies were highly consistent between replicates (Pearson’s pairwise correlation coefficient *r* between replicates: 0.946≤*r*≤ 0.999, Fig. S1). We used these frequencies to calculate the fitness of all DHFR variants relative to the wild type. Population genetic theory shows^47^ that it is best to represent all fitness values on a natural logarithmic scale (see Methods). On this scale, the fitness of the wild type has a value of 0, and that of a variant with relative fitness 1 corresponds to an exponential growth rate that is *e*^1^ ≈2.718 times higher.

The fitness values we measured are highly consistent with values obtained in a smaller independent experiment (*N*=250 variants, Pearson’s *r*=0.972, *p*=2×10^−158^, Fig. S2, A and B). Additional experiments confirmed that the genetic background of our *E. coli* strain did not substantially alter the relative fitness of our DHFR variants (see Methods and Fig. S2, C and D). Finally, to validate our fitness estimates, we isolated 30 DHFR variants and measuring their growth rate and resistance to trimethoprim in single cultures expressing individual DHFR variants. We found that relative fitness was highly correlated with the growth rate and resistance observed in single cultures (Fig. S2-4, Pearson’s *r*=0.993, *p*=2.3×10^−27^ for growth rate, Pearson’s *r*=0.987, *p*=1.1×10^−23^ for resistance).

### Functional DHFR variants vary widely in fitness and contain few amino acids at a key position

Most variants (93%, 243,303/261,332) have very low fitness (Fig. 1B). The distribution of their fitness values is consistent with that of variants with two stop codons (Fig. S5). Because DHFR catalytic activity bears a nearly linear relationship with *E. coli* fitness^48,49^, we conclude that DHFR is inactive in these variants. We thus also refer to these variants as *nonfunctional*, to distinguish them from the remaining 18,029 *functional* variants. The functional variants show highly variable fitness. Among them are high fitness variants identical to several previously characterized *folA* mutants^50^ with clinically relevant trimethoprim resistance levels that exceed wild type resistance by several orders of magnitude (Fig. S6). Almost all functional variants have a negatively charged aspartic or glutamic acid at position 27 (27Asp or 27Glu, Fig. S7A). This amino acid position is highly conserved in DHFR^51^ and highly prevalent in an alignment of 5000 orthologous DHFR proteins from proteobacteria (Fig. 1C). Consistent with a previous study that reported a functional mutant with a cysteine at position 27 (27Cys)^52^, we identified many functional 27Cys variants, even though the 27Cys allele is absent from the 5000 proteobacterial sequences we studied. We emphasize that the function-sequence relationship is not solely determined by position 27, for example because depending on the amino acid at position 27, different sets of amino acids are enriched at the neighboring positions 26 and 28 after selection (Fig. S8).

### The fitness landscape is rugged

To study the DHFR fitness landscape, we represented our data as a network or graph, in which each node (vertex) represents a DHFR variant, and is associated with the variant’s fitness. Any two DHFR variants that differ at one nucleotide position are immediate (1-mutant) neighbors connected by an edge, which represents a single mutational step. An evolutionary path through this network consists of several consecutive mutational steps. We restricted such paths to functional DHFR variants by removing all nonfunctional variants whose neighbors comprised only other nonfunctional variants. In contrast, we retained those nonfunctional variants with at least one functional neighbor in the network, reasoning that rare mutations may create a functional variant from a non-functional neighbor.

135,178 (52%) of DHFR variants in this network are contained in its largest connected subgraph (or “giant component”, refs.^53^), which constitutes the fitness landscape we analyze. Even though functional variants comprise only 13% of genotypes (18,019/135,178) of this landscape, they form a densely connected part of the landscape (Fig. S7, B and C), with as many functional-to-functional edges (50%, 161,015/324,044) as nonfunctional-to-functional edges. Almost 95% of functional variants had the maximally possible number of 27 (9×3) one-mutant neighbors, whereas nonfunctional variants had only one neighbor on average (Fig. S7, B and C).

Next we quantified our principal indicator of ruggedness, the number of peaks^8,10,13^, i.e. the number of DHFR variants that have higher fitness than all their one-mutant neighbors. We found that the landscape has 514 peaks and is thus rugged (Fig. 2A). In fact, a maximally rugged (uncorrelated) random NK landscape would contain a similar number of peaks^13^ (see Supplementary Note 1). Most of these peaks (408/514) have low fitness, i.e., they are less fit than the wild type (fitness < 0). 33 peaks have intermediate fitness (from 0 to 1) and were enriched with 27Cys variants. The remaining 73 peaks had high fitness (>1) and consisted exclusively of 27Asp and 27Glu variants (Fig. 2, A and B). Only one 27Asp peak has a fitness below 1, which equaled 0.87. For simplicity, we will refer to all 27Asp and 27Glu peaks (including this peak) as high fitness peaks.

**Fig. 2.**
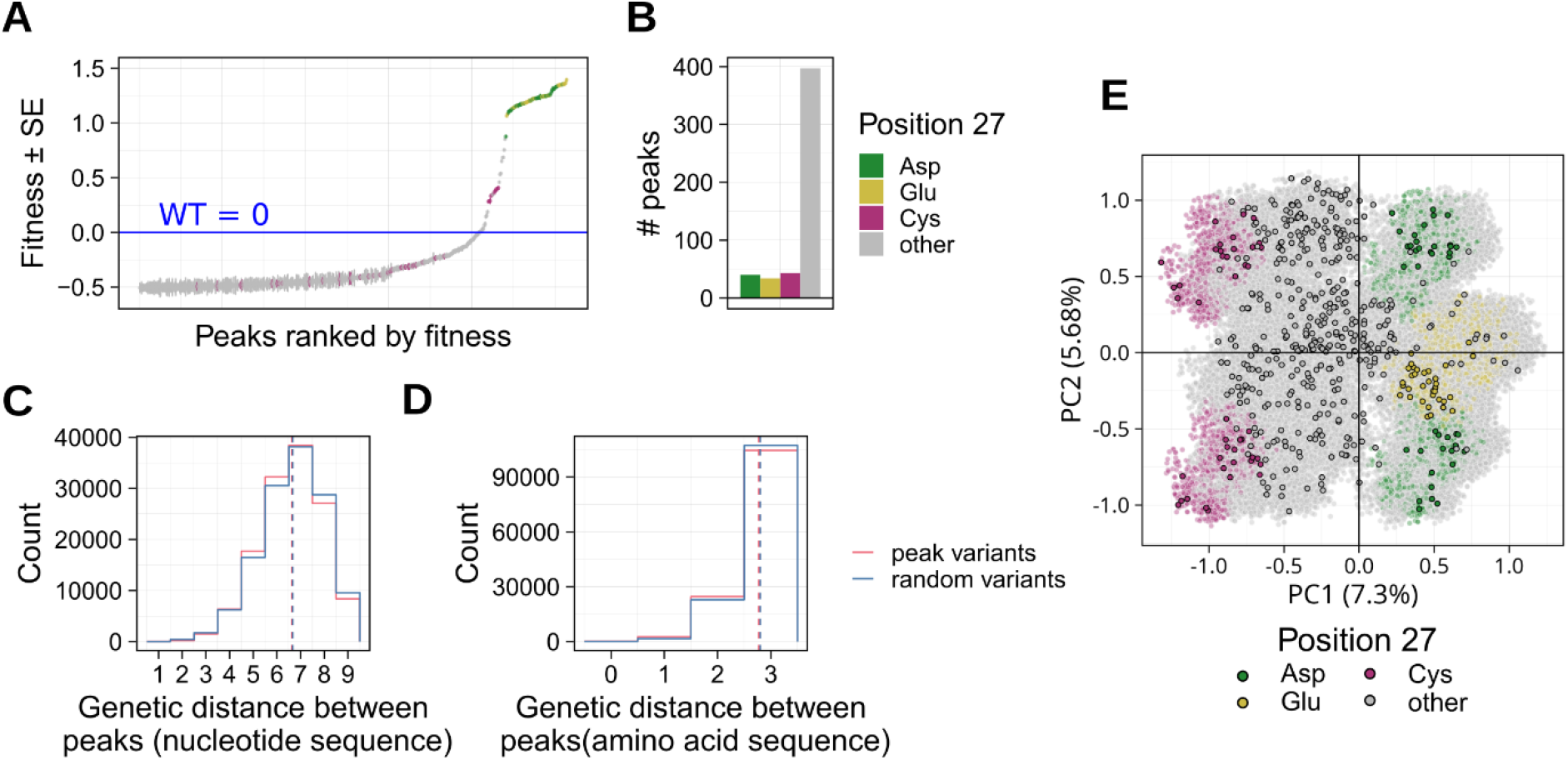
Fitness peaks. **(A) Multiple fitness peaks that vary widely in fitness**. The distribution of fitness estimates and corresponding standard errors for 514 variants identified as peaks in the landscape. The solid blue line at *y*=0 shows the fitness of the wild type. Colors indicate amino acid identity at position 27. **(B) Frequencies of amino acids at position 27 for fitness peaks** (*N*=514). **(C) Fitness peaks are genetically diverse**. Distribution of pairwise genetic distances between nucleotide sequences of 514 peaks (red) and between sequences of 514 randomly chosen variants (blue). The vertical lines show the means of the corresponding distributions. **(D)** As in panel **C** but based on amino acid sequences to calculate genetic distance. **(E) Principal component analysis shows that fitness peaks are spread across genotype space**. We used a matrix of pairwise genetic distances between all genotypes to perform the principal component analysis. The panel shows the projection on principal component 1 (PC1) and PC2. Each circle represents one of 135,178 variants in the landscape. Colors correspond to the amino acid at position 27. The circles with black outline are fitness peaks (*N*=514).

Because the location of different peaks in a landscape may affect their evolutionary accessibility, we next studied this location. Specifically, we asked whether peaks are close together in the landscape. We did so by computing the *genetic* (*nucleotide)* distance between all pairs of the 514 peaks, that is, the minimal number of single nucleotide changes that are needed to convert one peak into the other. We compared the resulting distance distribution with that of all pairs of 514 variants chosen at random from the landscape. Their mean distances are remarkably similar and lie within 1% percent of each other (*d*=6.67 for peaks vs. *d*=6.61 for random variants, two-sided Kolmogorov–Smirnov test *D* = 0.03, *p* < 10^−308^, *N* =131,841, Fig. 2C). This pattern also persists if we consider amino acid distances instead of nucleotide distances (Fig. 2D). Most importantly, the pattern extends to the high fitness peaks (i.e., 27Glu and 27Asp, Fig. S9). In sum, the DHFR landscape has many fitness peaks that are widely dispersed across the landscape (Fig. 2E).

### Fitness peaks are highly accessible

Considering the ruggedness of the landscape, one might expect that any one peak may only be accessible from a small fraction of genotypes. To find out, we first asked for each genotype and peak whether *accessible* paths exist from the genotype to the peak. Such paths consist only of beneficial (fitness-increasing) mutational steps, because fundamental population genetic principles dictate that weakly deleterious mutations are unlikely to go to fixation in an organism like *E.coli* with its large effective population size of 10^8^ individuals^54^.

More specifically, we determined the size of each peak’s *basin of attraction* – the total number of variants from which the peak is accessible. The size of this basin varied considerably with peak fitness, and with the amino acid at the critical position 27 (Fig. 3A). Low fitness peaks generally had small basins with a median size of only 28 variants (0.02% of all variants). Intermediate fitness peaks (27Cys) had larger basins with a median size of 7667 variants (5.7%). Most strikingly, high fitness peaks (with 27Glu and 27Asp) had very large basins whose median size comprised the majority (69% or 93,597) of variants. In general, peaks with higher fitness had significantly larger basins of attraction (Spearman’s ρ= 0.61, *p*=6.8×10^−55^, *N*=514, Fig. S10).

**Fig. 3.**
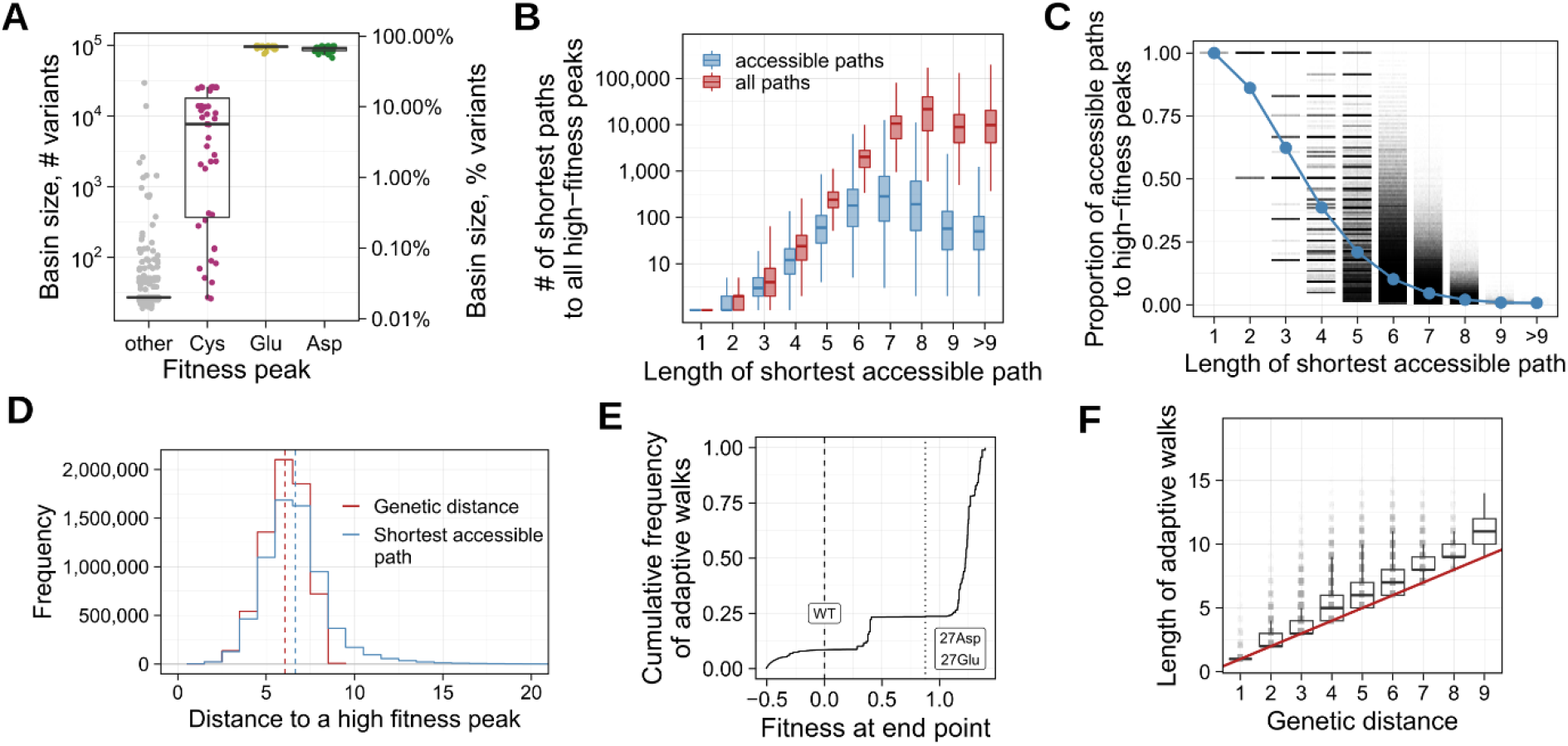
High fitness peaks are easily accessible. **(A) High fitness peaks have large basins of attraction.** The size of a peak’s basin of attraction depends on its amino acid at position 27 (color). Basin size is indicated both as the number of variants in the basin (left vertical axis), and as the percentage of the total number of the variants in the landscape (right vertical axis). **(B) The landscape contains many paths to high fitness peaks**. The vertical axis shows the total number of all shortest paths per variant to any high fitness peak. The horizontal axis shows the length of the shortest path. Red and blue boxplots summarize the number of paths for all shortest paths and all accessible shortest paths, respectively. Each box spans the interquartile range (IQR), each horizontal line inside a box indicates the median value, and each whisker extends to the minimum/maximum value outside the box within a 1.5 IQR interval. The data values beyond the 1.5 IQR interval is not shown. *N*=5,509,409,778 paths. **(C) The proportion of accessible paths depends on path length**. The vertical axis shows the proportion of accessible paths (out of all shortest paths) leading to high fitness peaks. As path length increases along the horizontal axis, the proportion of accessible paths decreases. Blue circles indicate the mean proportion of accessible paths at each length. *N*= 4,876,880 variant-peak pairs. **(D) Most accessible paths to fitness peaks are short**. The blue line shows the number of mutational steps for the shortest accessible paths leading from individual variants to each fitness peak accessible from this variant. The red line shows the distribution of the genetic distances for all corresponding variant-peak pairs (*N*=13,496,380). **(E) Most evolving populations reach high fitness peaks**. The panel shows the cumulative distribution of fitness values reached by 10^6^ adaptive walks starting from randomly selected variants in the landscape (Methods). The dashed vertical line *x*=0 shows the fitness value of the wild type. The dotted vertical line *x*=0.87 shows the fitness of the lowest among all high fitness peaks (27Asp/Glu). 76.5% of adaptive walks reached fitness 0.87 or higher. **(F) Most adaptive walks reaching high fitness peaks are short**. The vertical axis shows the length of adaptive walks (number of mutations) that started from a randomly selected variant and reached a high fitness peak. The horizontal axis shows the (shortest) genetic distances between each starting variant and the attained peak. The red line *y=x* shows the length of the shortest possible path to a high fitness peak, which is given by the genetic distance. The boxplots were constructed in the same way as in panel B. Adaptive walks were only marginally longer than shortest paths. *N*=765,181 adaptive walks.

The large basin size of the 27Asp/Glu peaks indicates their accessibility. However, even though accessible paths to any one high fitness peak may exist, they may be few compared to the total number of paths. To exclude this possibility, we focused on all variants within a peak’s basin of attraction and enumerated all shortest paths (accessible and inaccessible) from each variant to the peak. Not unexpectedly, the number of total paths increases exponentially with a variant’s distance from a peak (Fig. 3B). Among these paths, the fraction of *accessible* paths is very high at modest distances to a peak, but decreases with increasing path length (Fig. 3C). For example, at two, three, four, and five mutational steps away from a peak, 86%, 62%, 39%, and 21% of all paths are accessible, respectively. At the distance of 9 mutational steps, only 1% of paths remain accessible.

In a perfectly smooth landscape, the length of the shortest accessible path from any one variant to a high fitness peak equals the genetic distance between the variant and the peak. In contrast, in a rugged landscape even the shortest accessible path may meander through the landscape and thus be much longer than this genetic distance^50^. However, despite our landscape’s ruggedness, this is not the case. In our landscape, the mean length of shortest accessible paths between a variant and a high fitness peak (mean±s.d.=6.59±1.92 mutations) are less than one mutation longer than the mean genetic distance (5.99±1.25 mutations, Fig. 3D, two-sided Kolmogorov–Smirnov test *D*=0.15, *p*<10^−308^, *N*= 7,251,096).

### Adaptive evolution can easily reach high fitness peaks via short paths

Even though high fitness peaks appear highly accessible, their proportion (14%) is small compared to the majority of low fitness peaks. Would selection drive most evolving populations to one of the many (408) low fitness peaks? To find out, we simulated adaptive evolution in the weak-mutation strong-selection regime (SSWM, Methods)^55^. This choice is motivated by the extremely low mutation rate for the nine-nucleotide mutational target in the *E. coli* genome (= 9 positions × 2.2×10^−10^ substitutions per position per generation)^56^. In this regime, adaptive evolution effectively becomes an adaptive walk (Methods).

We simulated 10^6^ such walks, each starting from a randomly chosen DHFR variant. For all immediate neighbors that are only one mutational step away from this variant, we then calculated the fixation probability of the corresponding mutation. To this end, we used a well-established expression derived by Kimura for the probability that a new mutation sweeps through a population and becomes fixed^57^ (see Methods). This expression takes into account the fitness difference of the mutant to the starting variant, as well as the influence of genetic drift, whose strength falls with increasing effective population size. Because *E. coli* has very large populations (>10^8^ individuals), most fixation events are driven by selection. A population takes each mutational step with a probability that corresponds to this fixation probability. We repeated this procedure for every step of an adaptive walk until the walk reached a fitness peak.

Most adaptive walks reached a high fitness peak. Specifically, 76.5% of 10^6^ walks reached one of the 74 Glu/Asp high fitness peaks (Fig. 3E), corresponding to a trimethoprim resistance several orders of magnitudes higher than the resistance of the wild type (Fig. S6). A notable exception is the walks that started from 27Cys variants, 99.4% of which reached only 27Cys peaks with intermediate fitness (Fig. S11). Remarkably, peaks with higher fitness were not reached by fewer but by more adaptive walks (Spearman’s ρ= 0.62, *p*= 6.9×10^−55^, *N*=514, Fig. S12). Finally, adaptive walks terminating on high fitness peaks were relatively short, requiring only 5.6±2.1 mutational steps (mean±s.d.), or about one mutation more than the mean genetic distances between the corresponding start and end points of the adaptive walks (mean±s.d.=4.4±1.3, Fig. 3F, two-sided Kolmogorov–Smirnov test *D*=0.28, *p*<10^−308^, *N*=765,181).

Kimura’s fixation probability, which we used to model adaptive walks, favors the fixation of mutational steps with large fitness increases. This raises the possibility that the accessibility of the 27Glu/Asp peaks partly results from this procedure, because these peaks have very high fitness compared to the rest of the landscape. However, this is not the case. When we repeated simulations choosing each mutation to be fixed at random and uniformly among all fitness-increasing mutations, we found that 75% of walks still reached high fitness peaks (Fig. S13). This shows that the high accessibility of high fitness peaks is dictated by landscape topology rather than by details of the simulation procedure.

### Simultaneous accessibility of peaks leads to contingent evolution

Because high fitness peaks have enormous basins of attraction, individual variants must be members of multiple basins. To characterize this overlap between different basins quantitatively, we first determined the proportion of variants shared by any two basins. We found that 27Glu and 27Asp peaks shared 90.1±6.4% (mean ±s.d., *N*= 2,701) of the variants in their basins (Fig. 4A). In contrast, the basins of the 27Cys peaks shared a much smaller proportion of their variants on average (mean ±s.d.=22±29%, *N=*903). Furthermore, the basins of 27Glu/Asp peaks overlapped by only 1.6±1.6% (mean ±s.d., *N*= 3,182) of variants with those of 27Cys peaks (Fig. 4B). In other words, only a small proportion of variants had simultaneous access to both 27Glu/Asp and 27Cys peaks. Overall, 77% (104,496/135,176) of variants had access to more than one high fitness peak. Remarkably, 47.5% of all variants in the landscape had simultaneous access to all 74 high fitness peaks (Fig. 4C).

**Fig. 4.**
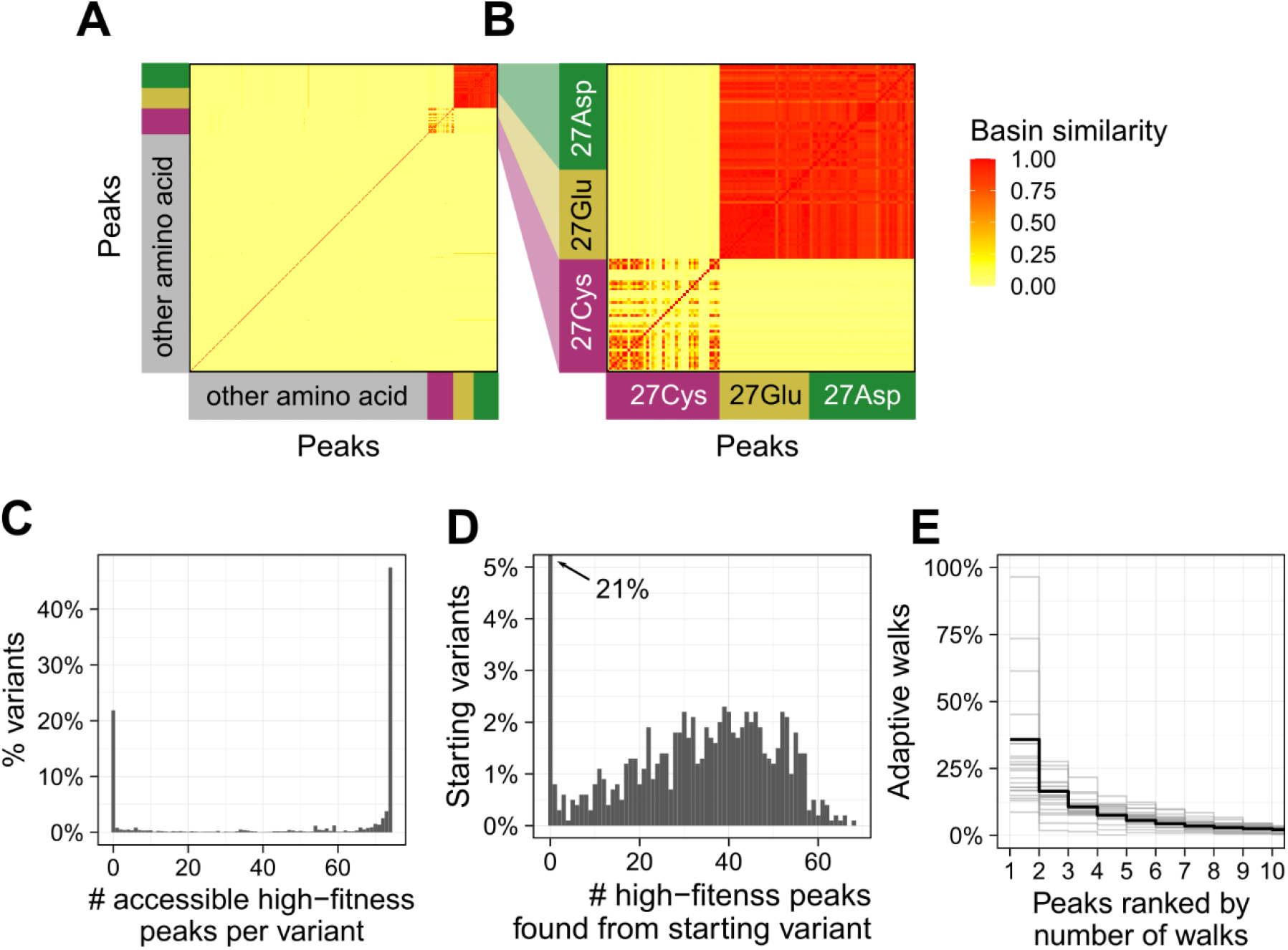
Evolutionary contingency. **(A) The basins of attraction of high fitness peaks overlap**. The heatmap shows the fraction of variants shared by different basins of attractions (see Methods). Each row and each column correspond to one peak, and peaks are classified according to the amino acid at position 27. A value of one (red) means that the basins of the corresponding peaks comprise identical sets of variants, and a value of zero (yellow) means that two basins do not share a single variant. The results are presented as a symmetric matrix of pairwise fractional overlaps between pairs of all 514 peaks. (**B)** A magnified portion of the matrix including only peaks containing 27Asp, 27Glu, and 27Cys. The basins of high fitness peaks (27Glu and 27 Asp) share more than 90% of their variants. **(C) Most variants have access to multiple high fitness peaks**. Distribution of the number of high fitness peaks (27Asp/Glu) that are accessible to each variant (*N*=135,178). **(D) Evolving populations starting from the same variant reach multiple peaks**. The distribution shows the number of high fitness peaks discovered by a total of 1000 populations evolving through adaptive walks that start from the same variant. Data is based on 10^6^ adaptive walks, such that 10^3^ walks started from the same starting variant among 10^3^ randomly chosen starting variants. Adaptive walks starting from 21% (209/1000) variants did not reach any high fitness peaks (vertical bar at *x*=0). For the remaining starting variants, adaptive walks reached multiple high fitness peaks. (**E) Adaptive walks preferentially attain some high fitness peaks**. The vertical axis shows the percentage of adaptive walks (out of 1000), which reached the most preferred peaks (horizontal axis). Each grey line summarizes the data for 1000 adaptive walks started from the same variant. To reduce visual clutter, grey lines are shown for only 24 randomly selected starting variants (24 grey lines) from which multiple high fitness peaks are reached. The black bold line shows the mean percentage for all 1000 starting variants.

Such simultaneous accessibility of multiple peaks can give rise to evolutionary contingency— the dependence of a historical process on chance events – because adaptive evolution starting from the same location can lead to different high fitness peaks. To test this hypothesis, we simulated 10^6^ further adaptive walks, such that 1000 walks started from each of 1000 randomly chosen starting variants. Adaptive walks originating from the same variant collectively reached 31 different high fitness peaks (mean±s.d.=29.5±17.8 peaks, median 31, Fig. 4D), confirming that simultaneous peak accessibility renders the identity of an attained fitness peak contingent on chance events during adaptive evolution.

During adaptive evolution, not all high fitness peaks are equally likely to be found by a population starting from a particular variant. For instance, 36±25% (mean±s.d.) of adaptive walks starting from the same location in the landscape reach a single, most commonly attained high fitness peak (Fig. 4E). But even walks converging to the same peak are likely to use different paths, because multiple paths to the same peak exist in its basin of attraction (24±57 mean±s.d. of shortest accessible paths from a variant to its peak, *N*=6,747,894, Fig. 3B). Adaptive walks indeed traverse such alternative paths (Fig. S14), which is another manifestation of contingency in our landscape. In sum, the large size and highly overlapping basins of attractions of different peaks render evolution on our DHFR landscape highly contingent on stochastic events.

## Discussion

The DHFR fitness landscape that we mapped through CRISPR-Cas9 gene editing is highly rugged, because it harbors more than 500 (mostly low) fitness peaks. At the same time, it has multiple properties expected from a smooth landscape. These include an abundance of monotonically fitness-increasing paths to high fitness peaks, enormous basins of attractions of these peaks, and easy reachability of these peaks by most evolving populations. High peak accessibility in a rugged landscape contradicts the predictions of classical computational models of random fitness landscapes, such as the NK landscape^15^. Biologically more realistic assumptions about the fitness distribution within a landscape would be required to explain our findings. For example, a recent study showed that a trade-off between fitness in environments with and without antibiotics can induce a smooth or rugged landscape, depending on the antibiotic concentration in the environment. An intermediate antibiotic concentration results in a landscape with the highest number of peaks, matching the conditions of our experiment. Although the high accessibility of fitness peaks and large basins of attraction of our landscape are also consistent with model predictions, our data does not satisfy the main assumption of this model, because it does not show a trade-off between fitness in the presence and absence of antibiotic (Fig S15). A biological realistic model to explain our observations thus remains an exciting task for future work.

Our metric of ruggedness – number of peaks – correlates with other such metrics^8,10,58^. Among the most widely used is the incidence of reciprocal sign epistasis (Fig. S16A), a kind of non-additive gene interaction that is necessary but not sufficient for the existence of multiple peaks^19,59,60^. We found that in the DHFR landscape 12.5% of mutant pairs show reciprocal sign epistasis (Fig. S16B). This value is within the range of other combinatorially complete landscapes (8-22%^39,58,61^), where peak accessibility has not been directly determined. In complex landscapes like these, the relationship between peak accessibility and reciprocal sign epistasis may be complex, and simple proxies of ruggedness may be less useful indicators of landscape navigability than is commonly assumed. Because reciprocal sign epistasis is not abundant, extradimensional by-passes^37,62,63^ – indirect paths that detour around a local fitness reduction caused by it – do also not play a major role in rendering fitness peaks accessible (Supplementary Note 2, Fig. S16, C and D)

Because *E.coli*’s effective population size exceeds 10^8^ individuals^54^, selection is much stronger than genetic drift in this species. For any organism with such large population size, even tiny differences in fitness are visible to natural selection. Even synonymous mutations can have a measurable effect on fitness, due to codon-specific effects on mRNA stability, and on the rate and fidelity of translation^64–67^. We thus considered all mutations (including synonymous mutations) in our landscape as non-neutral. However, we note that any existing technologies to measure fitness are limited by measurement error. Specifically, even in our six-fold replicated experiments and high sequencing coverage, we had limited power to detect fitness differences smaller than 2.5%. (The power to detect a 2.5% difference in fitness at *p=*0.05 equals *P=*0.933, Fig. S17). Such small fitness differences existed between 16% of all mutational neighbors in the DHFR landscape. If we had considered them effectively neutral, the number of accessible paths to high fitness peaks would have increased, because slightly deleterious mutational steps would have become accessible. In other words, assuming an absence of neutral mutations means that our high fitness peaks are even more accessible than we estimated – it renders our estimates of peak accessibility conservative.

The simultaneous accessibility of multiple fitness peaks results in evolutionary contingency at the level of the *genotype*, since different populations arrive at different high fitness peaks. Previous studies highlighted that genetic drift, mutational stochasticity, and epistasis can help create such contingency^50,68,69^. Our observations show that such contingency can even arise when drift is negligible. Importantly, our observations are consistent with previous work showing that genotypic contingency does not preclude the predictability of *phenotypic* evolution in a given environment^45,50,68,70^. For example, parallel populations from experimental evolution studies often evolve similar phenotypes via genetically distinct mutations^68,70^. Likewise, our analysis shows that adaptive evolution can reach genotypically different high fitness peaks with similar trimethoprim resistance.

However, genotypic contingency can entail phenotypic contingency and reduce the predictability of phenotypic evolution in a new environment^69^. This occurs whenever different genotypes with similar fitness in one environment have different fitness in another environment, creating bifurcation points at which evolutionary trajectories can diverge in a new environment^69^. Some of our high fitness genotypes illustrate this principle, because they differ in their fitness on a chemically modified version of trimethoprim^71^. While DHFR variants D27E and A26T are resistant against this new antibiotic, variant L28R is sensitive. All three substitutions occur in the high fitness peaks of our landscape. (A26T occurs in 15 peaks, D27E occurs in 34 peaks, and L28R occurs in one peak.) Depending on the peak they start from, a population subject to this new antibiotic would follow different evolutionary trajectories. More generally, because a change in environment may cause a deformation of fitness landscape helping to overcome low fitness valleys^72,73^ and because interactions between genotype and environment are frequent^74^, genotypic contingency may often entail phenotypic contingency.

Our landscape is one of the largest empirical landscapes currently available, but it covers a small segment of a single gene, whose variation constitutes a tiny fraction of sequence space. This is an unavoidable limitation of any empirical landscape study. It also makes generalization difficult because the choice of mutational target and sampling procedure can affect the properties of a reconstructed landscape^75^. We deliberately targeted a conservative gene that encodes a key metabolic enzyme in which a single mutation can lead to profound changes in fitness^48,76^. Consequently, only 9% of variants resulted in a functional enzyme.

Despite this constraint, functional variants formed a landscape with many fitness-increasing paths. A landscape derived from a less constrained target for gene editing would have harbored many more functional variants, and thus potentially even more paths accessible to Darwinian evolution than we observed. Future experiments using different mutational targets, organisms, sampling design, and experimental conditions will show whether very rugged yet highly navigable landscapes are typical or unusual.

When Sewall Wright coined the concept of an adaptive landscape nearly a century ago, he was concerned that multi-peaked landscapes may prevent adaptive Darwinian evolution driven by natural selection^1^. Simple theoretical models developed in the 20^th^ century support this concern^13,15^, but experimental evidence has been lacking. The landscape we studied shows that ruggedness need not impair Darwinian adaptation, even though it creates an enormous potential for contingent evolution. Our results suggest that we will need to refine our current theoretical understanding of the relationship between landscape ruggedness and navigability. Improved landscape theory will have to capture realistic properties of empirical landscapes, such as the sparsity of high-order epistasis^18^, the adaptational trade-offs between different fitness components^77^, strong local correlation in fitness among genotypes^40^, and dense connectivity of sequence space^37,78,79^. Because landscapes play an important role in fields as different as chemistry and the social sciences^2–7^, better data and theory on complex landscapes may also require other fields to re-evaluate the challenges of optimization problems on their landscapes. Even though a small fraction of peaks may have high fitness, in landscapes like that of DHFR these peaks can easily be discovered by simple evolutionary algorithms.

## Methods

Full methods are provided in the supplement

## Acknowledgements

This project has received funding from the European Research Council under Grant Agreement No. 739874. We would also like to acknowledge support by Swiss National Science Foundation grant 31003A_172887 and by the University Priority Research Program in Evolutionary Biology. A.P. is funded by a MSCA-IF-EF-ST 2020, 101030711.

LGP and JAE were supported by European Research Council Grant Agreement No. 803375 and the *Programa de Atracción de Talento de la Comunidad de Madrid* (grants 2016-T1/BIO-1105; and 2020-5A/BIO-19726). We thank to Diego Pesce and Caua Westmann for their help with establishing experimental protocols.

## Author contributions

A.P. and A.W conceived the study and designed experiments. A.P., L.G.P. and J. A. E. carried out experiments. A.P. and A.W. analyzed data. A.P. wrote computer code to carry out bioinformatic work, simulations and analysis. A.W. and A.P. wrote the paper, which was edited by all authors.

## References

1. Transactions and general addresses. in The roles of mutation, inbreeding, crossbreeding and selection in evolution (ed. Wright, S.) vol. 1 356–366 (Genetics Society of America, 1932).

2. Fear, K. K. & Price, T. The Adaptive Surface in Ecology. Oikos 82, 440–448 (1998).

3. Stadler, B. M. R. & Stadler, P. F. Generalized Topological Spaces in Evolutionary Theory and Combinatorial Chemistry. J. Chem. Inf. Comput. Sci. 42, 577–585 (2002).

4. Mühlenbein, H. Evolution in time and space - the parallel genetic algorithm. In Foundations of Genetic Algorithms 316–337 (Morgan Kaufmann, 1991).

5. Manderick, B., Weger, M. K. de & Spiessens, P. The Genetic Algorithm and the Structure of the Fitness Landscape. in Proceedings of the 4th International Conference on Genetic Algorithms, San Diego, CA, USA, July 1991 (eds. Belew, R. K. & Booker, L. B.) 143–150 (Morgan Kaufmann, 1991).

6. Levinthal, D. A. Organizational Adaptation and Environmental Selection-Interrelated Processes of Change. Organ. Sci. 2, 140–145 (1991).

7. Vassilev, V. K., Muller, J. F. & Fogarty, T. C. Digital circuit evolution and fitness landscapes. in Proceedings of the 1999 Congress on Evolutionary Computation-CEC99 (Cat. No. 99TH8406) vol. 2 1299–1306 Vol. 2 (1999).

8. de Visser, J. A. G. M. & Krug, J. Empirical fitness landscapes and the predictability of evolution. Nat. Rev. Genet. 15, 480–490 (2014).

9. Fragata, I., Blanckaert, A., Dias Louro, M. A., Liberles, D. A. & Bank, C. Evolution in the light of fitness landscape theory. Trends Ecol. Evol. 34, 69–82 (2019).

10. Szendro, I. G., Schenk, M. F., Franke, J., Krug, J. & Visser, J. A. G. M. de. Quantitative analyses of empirical fitness landscapes. J. Stat. Mech. Theory Exp. 2013, P01005 (2013).

11. Weinreich, D. M., Watson, R. A. & Chao, L. Perspective: Sign Epistasis and Genetic Costraint on Evolutionary Trajectories. Evolution 59, 1165–1174 (2005).

12. Bank, C. Epistasis and Adaptation on Fitness Landscapes. Annu. Rev. Ecol. Evol. Syst. 53, ull (2022).

13. Kauffman, S. & Levin, S. Towards a general theory of adaptive walks on rugged landscapes. J. Theor. Biol. 128, 11–45 (1987).

14. Hegarty, P. & Martinsson, A. On the existence of accessible paths in various models of fitness landscapes. Ann. Appl. Probab. 24, (2014).

15. Kauffman, S. A. & Weinberger, E. D. The NK model of rugged fitness landscapes and its application to maturation of the immune response. J. Theor. Biol. 141, 211–245 (1989).

16. Kingman, J. F. C. A simple model for the balance between selection and mutation. J. Appl. Probab. 15, 1–12 (1978).

17. Aita, T. & Husimi, Y. Fitness Spectrum Among Random Mutants on Mt. Fuji-Type Fitness Landscape. J. Theor. Biol. 182, 469–485 (1996).

18. Poelwijk, F. J., Socolich, M. & Ranganathan, R. Learning the pattern of epistasis linking genotype and phenotype in a protein. Nat. Commun. 10, 1–11 (2019).

19. Crona, K., Greene, D. & Barlow, M. The peaks and geometry of fitness landscapes. J. Theor. Biol. 317, 1–10 (2013).

20. Lozovsky, E. R. et al. Stepwise acquisition of pyrimethamine resistance in the malaria parasite. Proc. Natl. Acad. Sci. 106, 12025–12030 (2009).

21. Khan, A. I., Dinh, D. M., Schneider, D., Lenski, R. E. & Cooper, T. F. Negative Epistasis Between Beneficial Mutations in an Evolving Bacterial Population. Science 332, 1193– 1196 (2011).

22. O’Maille, P. E. et al. Quantitative exploration of the catalytic landscape separating divergent plant sesquiterpene synthases. Nat. Chem. Biol. 4, 617–623 (2008).

23. Hall, D. W., Agan, M. & Pope, S. C. Fitness Epistasis among 6 Biosynthetic Loci in the Budding Yeast Saccharomyces cerevisiae. J. Hered. 101, S75–S84 (2010).

24. da Silva, J., Coetzer, M., Nedellec, R., Pastore, C. & Mosier, D. E. Fitness Epistasis and Constraints on Adaptation in a Human Immunodeficiency Virus Type 1 Protein Region. Genetics 185, 293–303 (2010).

25. Tan, L., Serene, S., Chao, H. X. & Gore, J. Hidden Randomness between Fitness Landscapes Limits Reverse Evolution. Phys. Rev. Lett. 106, 198102 (2011).

26. Lunzer, M., Miller, S. P., Felsheim, R. & Dean, A. M. The Biochemical Architecture of an Ancient Adaptive Landscape. Science 310, 499–501 (2005).

27. Reetz, M. T. & Sanchis, J. Constructing and Analyzing the Fitness Landscape of an Experimental Evolutionary Process. ChemBioChem 9, 2260–2267 (2008).

28. Brown, K. M. et al. Compensatory Mutations Restore Fitness during the Evolution of Dihydrofolate Reductase. Mol. Biol. Evol. 27, 2682–2690 (2010).

29. Chou, H.-H., Chiu, H.-C., Delaney, N. F., Segrè, D. & Marx, C. J. Diminishing Returns Epistasis Among Beneficial Mutations Decelerates Adaptation. Science 332, 1190–1192 (2011).

30. Bank, C., Matuszewski, S., Hietpas, R. T. & Jensen, J. D. On the (un)predictability of a large intragenic fitness landscape. Proc. Natl. Acad. Sci. 113, 14085–14090 (2016).

31. Olson, C. A., Wu, N. C. & Sun, R. A Comprehensive Biophysical Description of Pairwise Epistasis throughout an Entire Protein Domain. Curr. Biol. 24, 2643–2651 (2014).

32. Sarkisyan, K. S. et al. Local fitness landscape of the green fluorescent protein. Nature 533, 397–401 (2016).

33. Rowe, W. et al. Analysis of a complete DNA-protein affinity landscape. J. R. Soc. Interface 7, 397–408 (2010).

34. Warren, C. L. et al. Defining the sequence-recognition profile of DNA-binding molecules. Proc. Natl. Acad. Sci. 103, 867–872 (2006).

35. Payne, J. L. & Wagner, A. The causes of evolvability and their evolution. Nat. Rev. Genet. 20, 24–38 (2019).

36. Jiménez, J. I., Xulvi-Brunet, R., Campbell, G. W., Turk-MacLeod, R. & Chen, I. A. Comprehensive experimental fitness landscape and evolutionary network for small RNA. Proc. Natl. Acad. Sci. U. S. A. 110, 14984–14989 (2013).

37. Wu, N. C., Dai, L., Olson, C. A., Lloyd-Smith, J. O. & Sun, R. Adaptation in protein fitness landscapes is facilitated by indirect paths. eLife 5, e16965 (2016).

38. Pokusaeva, V. O. et al. An experimental assay of the interactions of amino acids from orthologous sequences shaping a complex fitness landscape. PLOS Genet. 15, e1008079 (2019).

39. Domingo, J., Diss, G. & Lehner, B. Pairwise and higher-order genetic interactions during the evolution of a tRNA. Nature 558, 117–121 (2018).

40. Kuo, S.-T. et al. Global fitness landscapes of the Shine-Dalgarno sequence. Genome Res. 30, 711–723 (2020).

41. Faure, A. J. et al. Mapping the energetic and allosteric landscapes of protein binding domains. Nature 604, 175–183 (2022).

42. Lite, T.-L. V. et al. Uncovering the basis of protein-protein interaction specificity with a combinatorially complete library. eLife 9, e60924 (2020).

43. Aakre, C. D. et al. Evolving New Protein-Protein Interaction Specificity through Promiscuous Intermediates. Cell 163, 594–606 (2015).

44. Reisch, C. R. & Prather, K. L. J. The no-SCAR (Scarless Cas9 Assisted Recombineering) system for genome editing in Escherichia coli. Sci. Rep. 5, (2015).

45. Toprak, E. et al. Evolutionary paths to antibiotic resistance under dynamically sustained drug selection. Nat. Genet. 44, 101–105 (2011).

46. Tamer, Y. T. et al. High-Order Epistasis in Catalytic Power of Dihydrofolate Reductase Gives Rise to a Rugged Fitness Landscape in the Presence of Trimethoprim Selection. Mol. Biol. Evol. 36, 1533–1550 (2019).

47. Crow, J. F. & Kimura, M. An introduction to population genetics theory. (Scientific Publisher (India) ; The Blackburn Press, 2010).

48. Thompson, S., Zhang, Y., Ingle, C., Reynolds, K. A. & Kortemme, T. Altered expression of a quality control protease in E. coli reshapes the in vivo mutational landscape of a model enzyme. eLife 9, e53476 (2020).

49. Rodrigues, J. V. et al. Biophysical principles predict fitness landscapes of drug resistance. Proc. Natl. Acad. Sci. U. S. A. 113, E1470–1478 (2016).

50. Palmer, A. C. et al. Delayed commitment to evolutionary fate in antibiotic resistance fitness landscapes. Nat. Commun. 6, 1–8 (2015).

51. Volpato, J. P. & Pelletier, J. N. Mutational ‘hot-spots’ in mammalian, bacterial and protozoal dihydrofolate reductases associated with antifolate resistance: Sequence and structural comparison. Drug Resist. Updat. 12, 28–41 (2009).

52. Rodrigues, J. V. & Shakhnovich, E. I. Adaptation to mutational inactivation of an essential gene converges to an accessible suboptimal fitness peak. eLife 8, e50509 (2019).

53. Clark, J. A first look at graph theory. (World Scientific, 2005).

54. Lynch, M. et al. Genetic drift, selection and the evolution of the mutation rate. Nat. Rev. Genet. 17, 704–714 (2016).

55. Gillespie, J. H. Some Properties of Finite Populations Experiencing Strong Selection and Weak Mutation. Am. Nat. 121, 691–708 (1983).

56. Lee, H., Popodi, E., Tang, H. & Foster, P. L. Rate and molecular spectrum of spontaneous mutations in the bacterium Escherichia coli as determined by whole-genome sequencing. Proc. Natl. Acad. Sci. U. S. A. 109, E2774–2783 (2012).

57. Kimura, M. On the Probability of Fixation of Mutant Genes in a Population. Genetics 47, 713–719 (1962).

58. Song, S. & Zhang, J. Unbiased inference of the fitness landscape ruggedness from imprecise fitness estimates. Evolution 75, 2658–2671 (2021).

59. Poelwijk, F. J., Tănase-Nicola, S., Kiviet, D. J. & Tans, S. J. Reciprocal sign epistasis is a necessary condition for multi-peaked fitness landscapes. J. Theor. Biol. 272, 141–144 (2011).

60. Riehl, M., Phillips, R., Pudwell, L. & Chenette, N. Occurrences of reciprocal sign epistasis in single- and multi-peaked theoretical fitness landscapes. J. Phys. Math. Theor. 55, 434002 (2022).

61. Li, C., Qian, W., Maclean, C. J. & Zhang, J. The fitness landscape of a tRNA gene. Science 352, 837–840 (2016).

62. Conrad, M. The geometry of evolution. Biosystems 24, 61–81 (1990).

63. Chen, J.-C. & Conrad, M. A multilevel neuromolecular architecture that uses the extradimensional bypass principle to facilitate evolutionary learning. Phys. Nonlinear Phenom. 75, 417–437 (1994).

64. Boël, G. et al. Codon influence on protein expression in E. coli correlates with mRNA levels. Nature 529, 358–363 (2016).

65. Plotkin, J. B. & Kudla, G. Synonymous but not the same: the causes and consequences of codon bias. Nat. Rev. Genet. 12, 32–42 (2011).

66. Drummond, D. A. & Wilke, C. O. Mistranslation-Induced Protein Misfolding as a Dominant Constraint on Coding-Sequence Evolution. Cell 134, 341–352 (2008).

67. Shen, X., Song, S., Li, C. & Zhang, J. Synonymous mutations in representative yeast genes are mostly strongly non-neutral. Nature 606, 725–731 (2022).

68. Blount, Z. D., Lenski, R. E. & Losos, J. B. Contingency and determinism in evolution: Replaying life’s tape. Science 362, eaam5979 (2018).

69. Blount, Z. D., Borland, C. Z. & Lenski, R. E. Historical contingency and the evolution of a key innovation in an experimental population of Escherichia coli. Proc. Natl. Acad. Sci. U. S. A. 105, 7899–7906 (2008).

70. Meyer, J. R. et al. Repeatability and Contingency in the Evolution of a Key Innovation in Phage Lambda. Science 335, 428–432 (2012).

71. Manna, M. S. et al. A trimethoprim derivative impedes antibiotic resistance evolution. Nat. Commun. 12, 2949 (2021).

72. Mustonen, V. & Lässig, M. From fitness landscapes to seascapes: non-equilibrium dynamics of selection and adaptation. Trends Genet. 25, 111–119 (2009).

73. Steinberg, B. & Ostermeier, M. Environmental changes bridge evolutionary valleys. Sci. Adv. 2, e1500921 (2016).

74. Li, C. & Zhang, J. Multi-environment fitness landscapes of a tRNA gene. Nat. Ecol. Evol. 2, 1025–1032 (2018).

75. Whitlock, M. C., Phillips, P. C., Moore, F. B.-G. & Tonsor, S. J. Multiple Fitness Peaks and Epistasis. Annu. Rev. Ecol. Syst. 26, 601–629 (1995).

76. Bershtein, S., Choi, J.-M., Bhattacharyya, S., Budnik, B. & Shakhnovich, E. Systems-Level Response to Point Mutations in a Core Metabolic Enzyme Modulates Genotype-Phenotype Relationship. Cell Rep. 11, 645–656 (2015).

77. Das, S. G., Direito, S. O., Waclaw, B., Allen, R. J. & Krug, J. Predictable properties of fitness landscapes induced by adaptational tradeoffs. eLife 9, e55155 (2020).

78. Franke, J., Klözer, A., Visser, J. A. G. M. de & Krug, J. Evolutionary Accessibility of Mutational Pathways. PLOS Comput. Biol. 7, e1002134 (2011).

79. Greenbury, S. F., Louis, A. A. & Ahnert, S. E. The structure of genotype-phenotype maps makes fitness landscapes navigable. Nat. Ecol. Evol. 6, 1742–1752 (2022).

