## Supplementary materials for "A rugged yet easily navigable fitness landscape of antibiotic resistance"

Supplementary Figures

Supplementary Tables

Supplementary Methods

Supplementary Notes

Supplementary References

### Supplementary Figures

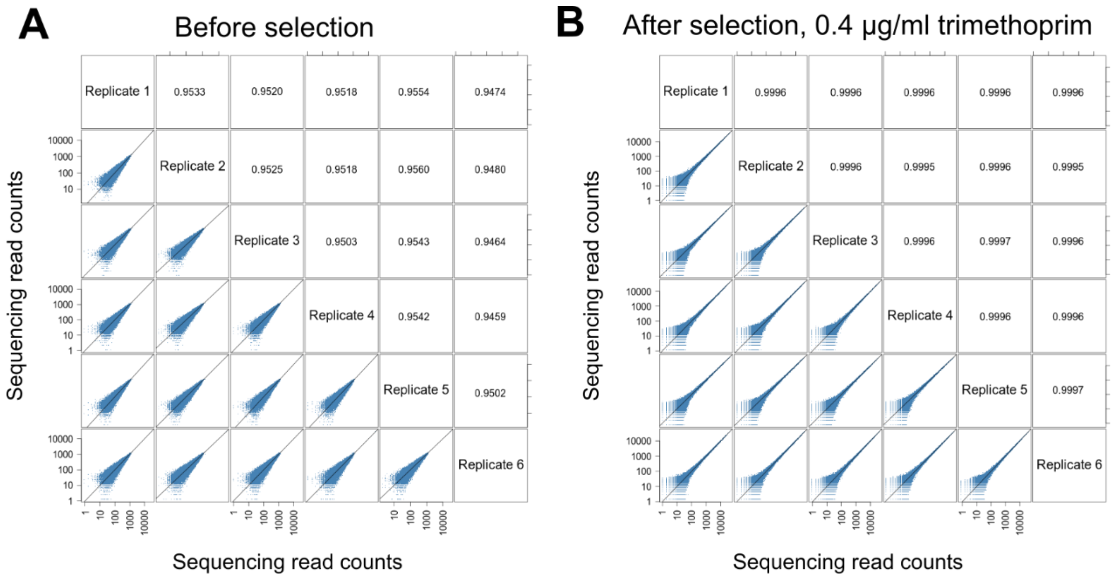


**Fig. S1. Correlation of read coverage in the *folA* mutant library** between replicates **(A)** before selection and **(B)** after selection**.** Note the logarithmic scale in all panels.

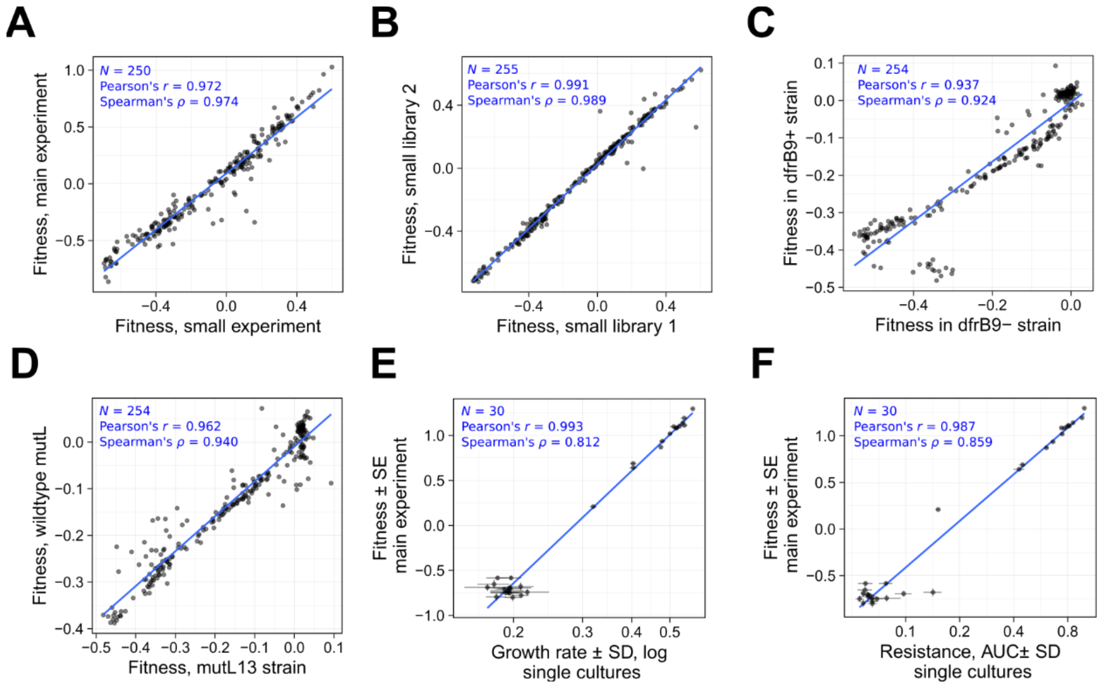


**Fig. S2. Reproducibility of mass selection experiment. (A)** Correlation in fitness between main mass selection experiment and a smaller independent experiment. **(B)** Correlation in fitness between two (out of six) small libraries, created by gene editing independently and selected in parallel. **(C)** Correlation in fitness of *folA* mutants obtained in the *E.coli* strain with dfrB9 integration and without dfrB9 integration. **(D)** Correlation in fitness of *folA* mutants obtained in the mutator strain (genotype mutL13) and in the wildtype *mutL* genetic background. **(E)** Correlation between fitness measured using the main selection experiment and the maximal growth rate estimated in single cultures. We isolated 30 clones from the library and grew single cultures (4 replicates per clone) to obtain growth curves with 0.4 µg/ml trimethoprim (shown in Fig. S3). We determined the maximal growth rate during exponential growth as the number of doublings per hour. The horizontal error bars show the standard deviation of growth rate for four replicated cultures. The vertical error bars show the standard errors for a maximum likelihood estimate of relative fitness (see Methods). **(F)** Correlation between fitness measured using the main selection experiment and the trimethoprim resistance for the same set of 30 clones as in panel E. We incubated single cultures at ten trimethoprim concentrations (from 0.0625 to 32 µg /ml) and used end-point optical densities after 24 h to obtain dose-response curves (shown on Fig. S4). We calculated the area under the dose-response curves (AUC, see Methods) and used it as a measure of resistance. The horizontal errors show the standard deviation of AUC for four replicates of the same clone. The vertical errors show the standard errors for a maximum likelihood estimate of relative fitness (see Methods).

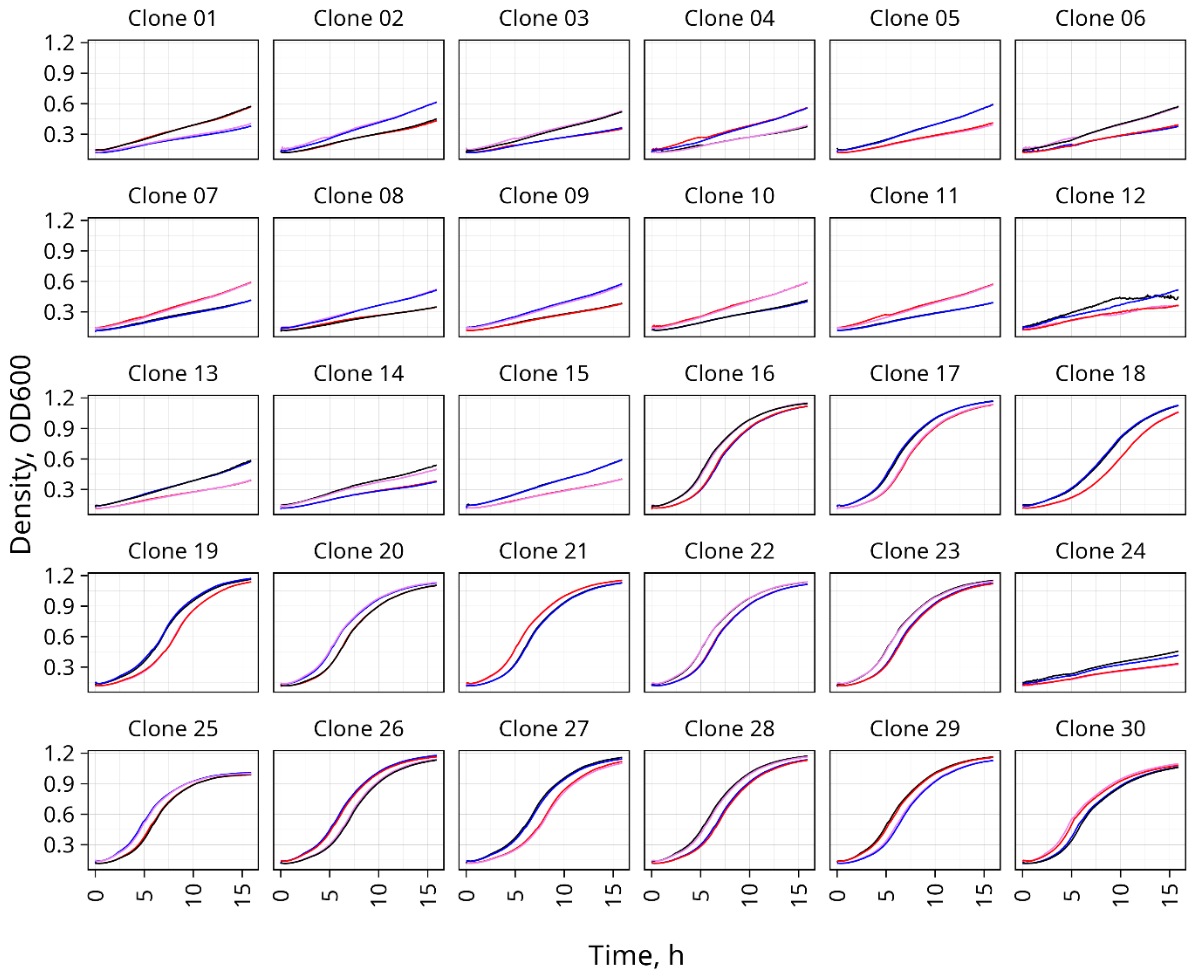


**Fig. S3. Growth curves of 30 variants in single cultures.** The panels show the change in the optical density ($OD_{600}$, the vertical axis) in M9 medium with 0.4 µg/ml trimethoprim over 16 hours (horizontal axis). Each panel shows growth curves for one of the 30 isolated clones. We measured the growth curve of each clone using 4 replicates (indicated by different colors in each panel. More specifically, we grouped the four replicates into two blocks, and measured the growth of each two-block group on different days (two replicates per block). Due to the small variation within each block, each two replicates appear almost identical. We used the growth curves to estimate the maximal growth rate shown in Fig. S2E (see Methods).

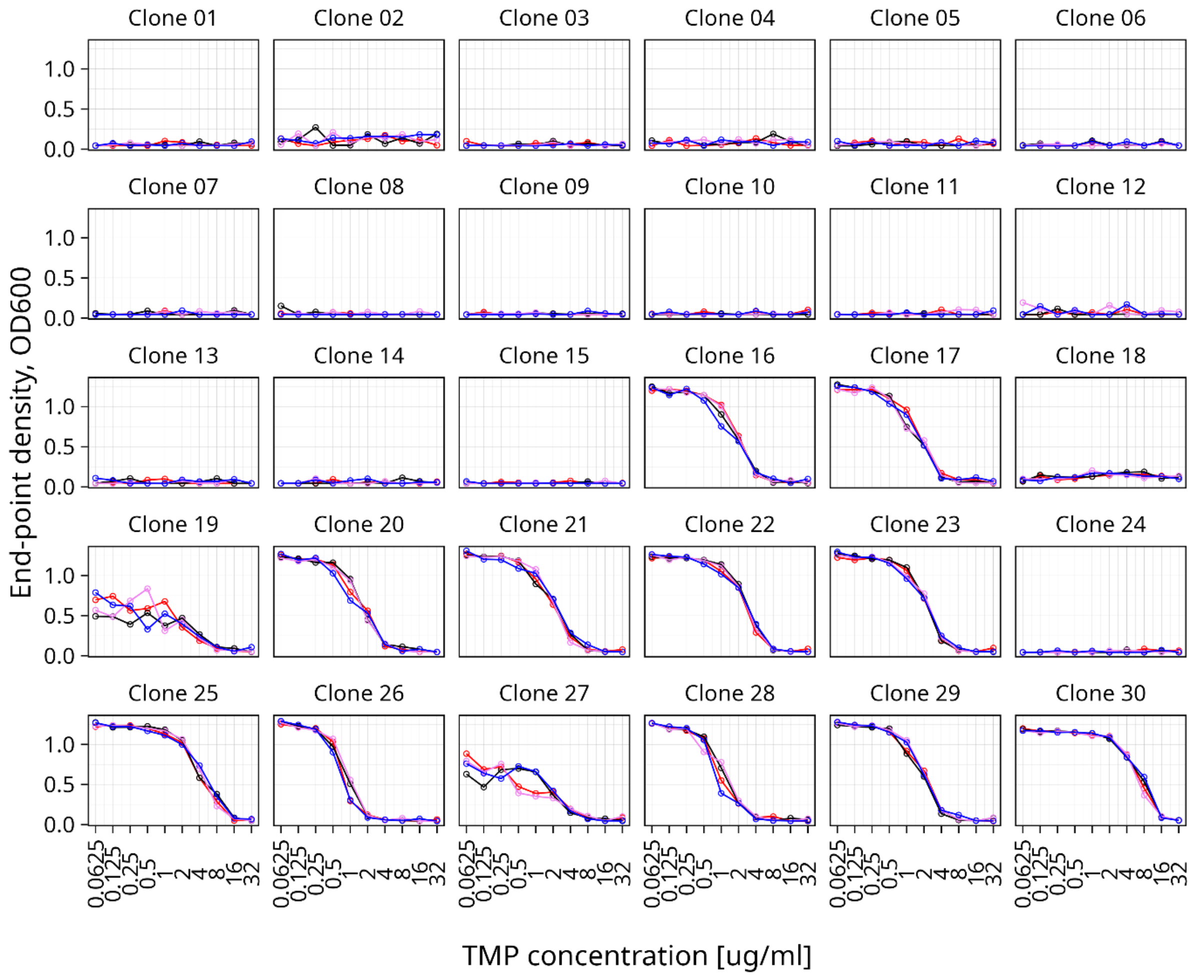


**Fig. S4. Trimethoprim resistance of 30 DHFR variants.** The vertical axis shows end-point optical density ($OD_{600}$) in single cultures incubated at different concentrations of trimethoprim (horizontal axis) after 24 h. Each panel shows dose-response curves for one of the 30 variants with 4 replicates per variant (indicated by color). We used the area under the dose response curves (AUC) as a measure of resistance (see Fig. S2F and Methods).

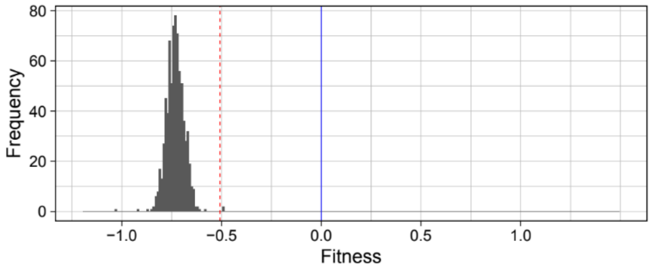


**Fig. S5. Distribution of fitness effects of DHFR variants with two stop codons (*N*=752).** We measured the fitness of double nonsense mutants relative to the wild type (blue line, fitness=0). The red dotted line shows the cutoff for nonfunctional variants (fitness = -0.508), estimated as explained in Fig. 1B and in Methods.

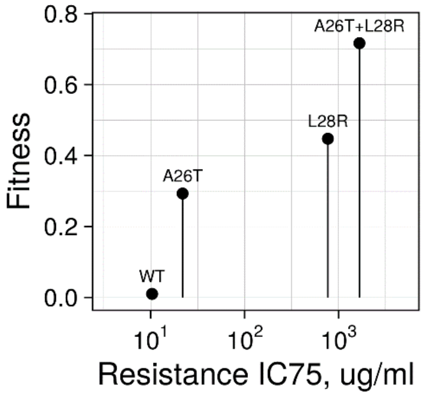


**Fig. S6. The resistance of previously characterized mutations and their fitness in our mass competition experiment.** We compare the fitness of three previously characterized DHFR mutations (A26T, L28R, and the double mutant A26T+L28R, Palmer et al. 2015) and the wild type. The horizontal axis shows trimethoprim resistance as inhibitory concentration IC75 from a previous study[50]. The vertical axis shows the relative fitness values from our main mass selection experiment. Please note that Palmer et al. found higher trimethoprim resistance compared to what we observed in our isolates (Fig. S4). This can be explained by the lower growth rate of our parental strain in comparison to the wild type MG1655 *E. coli* (Fig. S24). Moreover, Palmer et al. determined resistance after 36 h of incubation, whereas we used 24 h of incubation.

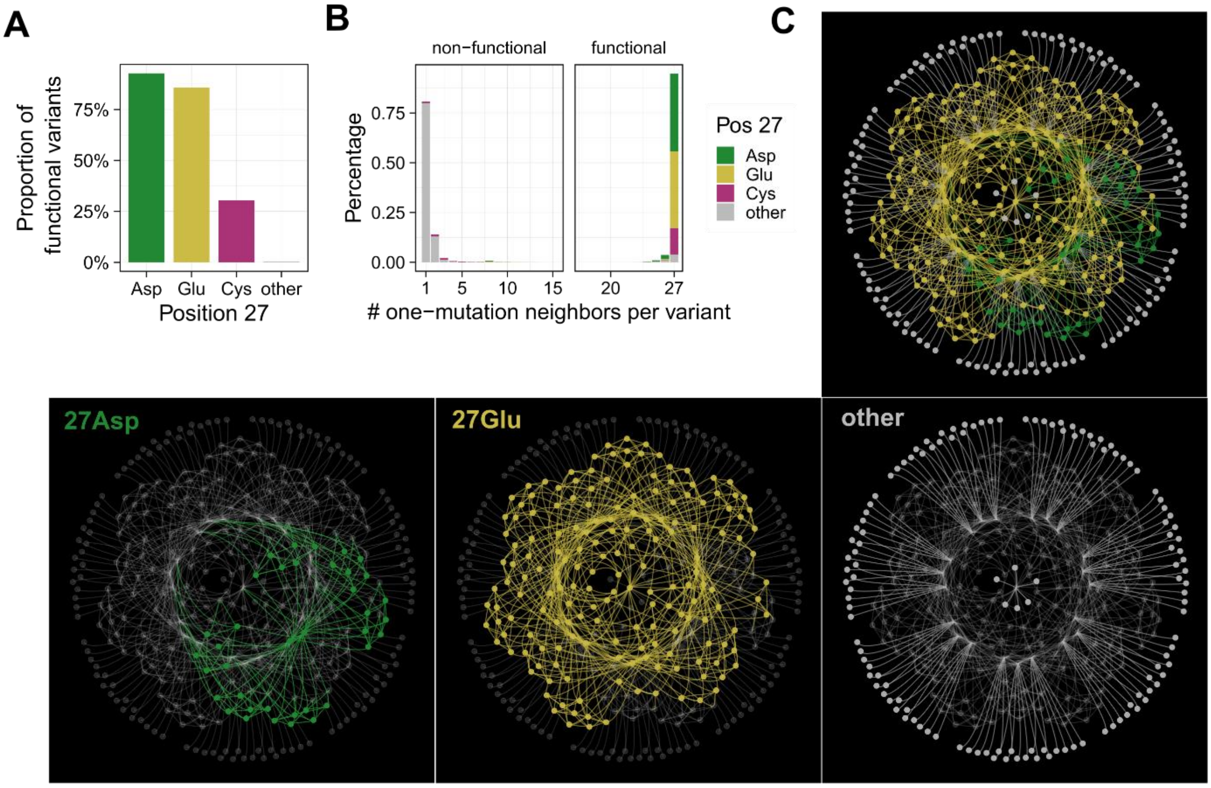


**Fig. S7. Local structure of the landscape. (A) Three amino acids at DHFR position 27 produce most functional variants.** The proportion of functional variants in the library, as a function of the amino acid at position 27. **(B) Functional variants are highly connected in the landscape.** The number of one-mutation neighbors per variant in the landscape, depending on the amino acid at position 27 (color legend), for non-functional variants (left panel) and for functional variants (right panel). Note that functional variants are connected to multiple neighbors, whereas non-functional variants are connected to only one neighbor. **(C) The structure of the neighborhood for a representative functional variant.** The four panels show the same subgraph for the neighborhood of a representative 27Glu variant ("AACGATACT"). The subgraph includes this variant at the center, surrounded by all other variants that differ by no more than two mutations from the focal variant. The color of nodes (variants) in the subgraphs correspond to the amino acid at position 27 (yellow: 27Glu; green: 27Asp; grey: all other amino acids). The top panel shows all variants in the subgraph. In the bottom panels, one of the amino acid 27 genotypes of variants is highlighted (27Glu, 27Asp, other amino acids) and the other types are not shown. All 27Glu and 27Asp variants in this subgraph are functional ($N$=208) and all variants with other amino acids at position 27 are non-functional ($N$=134). This subgraph shows that functional variants have many neighbors, whereas nonfunctional variants have one or few neighbors (see also panel **B**). The neighborhood structure of other functional variants is similar to that shown here.

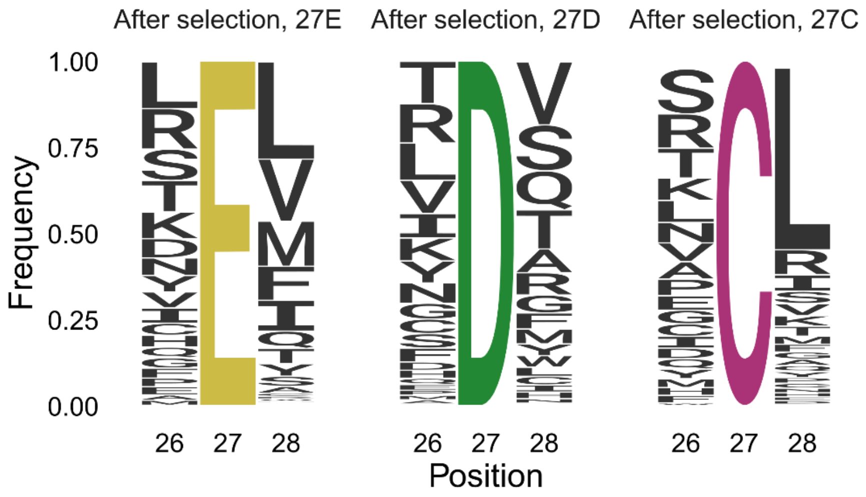


**Fig. S8. Amino acid composition after mass selection at position 26 and 28 depends on the amino acid at position 27.** Mass selection of variants with amino acid identity E, D or C at position 27 resulted in enrichment of different amino acids at neighboring positions 26 and 28. Compare, for example, the ranking of S at position 26 or that of L at position 28.

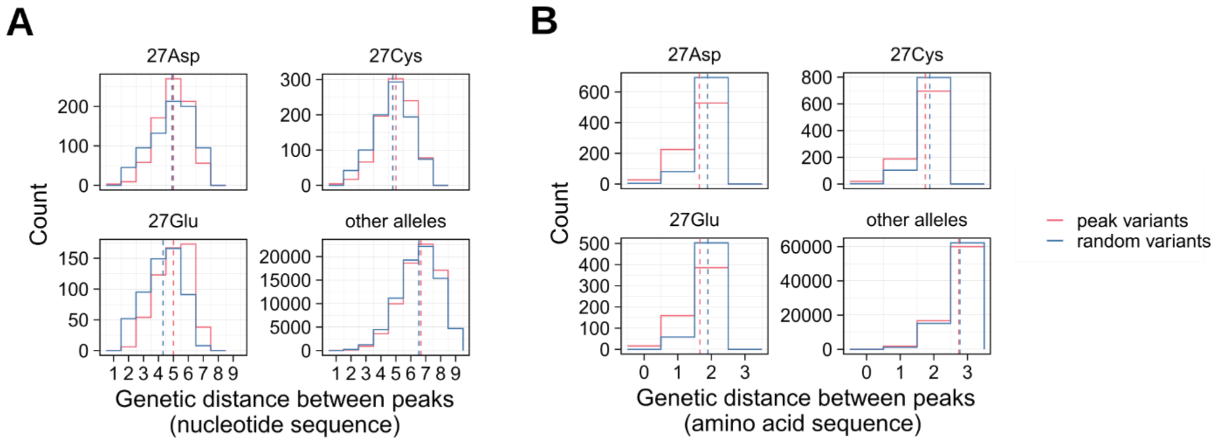


**Fig. S9. Distance between fitness peaks. (A)** Distribution of pairwise genetic distance between peaks (red) or between random pairs of variants (blue). We restricted the distance calculation for each panel to a subset of peaks or variants in one of the four allele classes at DHFR position 27 (Asp, Glu, Cys, other amino acids). We used the same number of random variants as the number of peaks. Vertical lines show mean distance values for the peaks (red) and the random variants (blue). **(B)** The same analysis of pairwise distances as in panel A, but using amino acid sequence instead of nucleotide sequence.

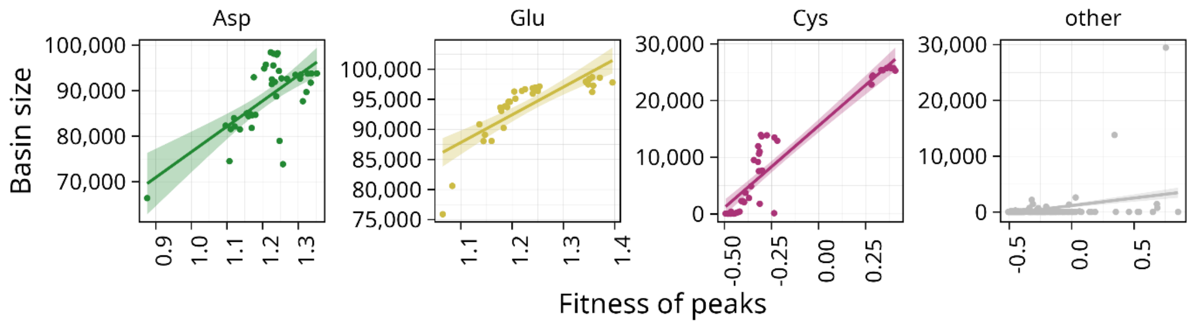


**Fig. S10. Peaks with higher fitness (horizontal axis) have larger basins of attraction (vertical axis).** Panels group peaks depending on the allele at position 27 (Asp, Glu, Cys, other amino acids). Circles show data for individual peaks, straight lines show model fits, and shaded regions show standard errors based on a linear model (Spearman's rank correlation for Asp peaks $\rho = 0.53$, $p=5.29\times10^{-4}$, $N=40$; for Glu peaks $\rho = 0.92$, $p<10^{-16}$, $N=34$; for Cys peaks $\rho = 0.92$, $p<10^{-16}$, $N=43$; for other peaks $\rho = 0.28$, $p=9.02\times10^{-9}$, $N=397$).

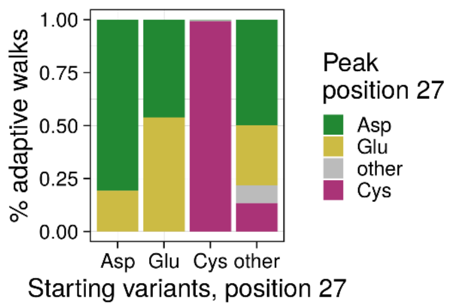


**Fig. S11. The genotype of a starting point in an adaptive walk influences the end genotype.** The distribution of the amino acids at key position 27 (vertical axis, color legend) of those peak variants that were reached from starting variants with different amino acids at position 27 (horizontal axis). The panel summarizes results for $10^6$ adaptive walks.

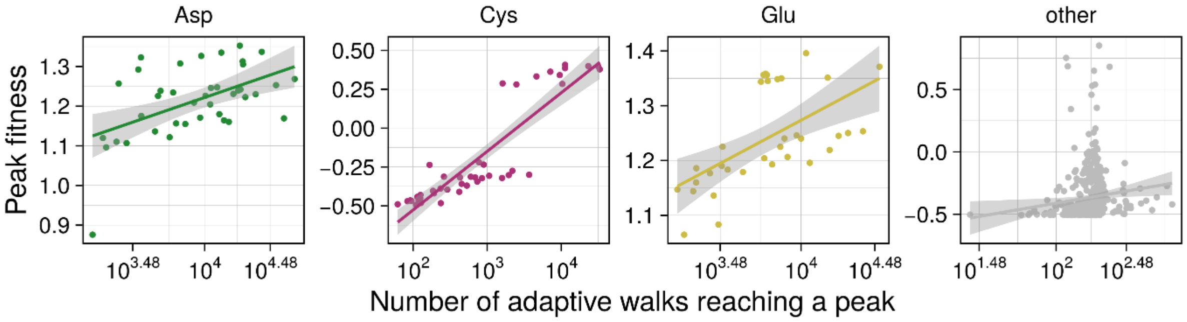


**Fig. S12. Peaks with higher fitness (vertical axis) are reached by more adaptive walks (horizontal axis).** Panels group peaks depending on the allele at position 27 (Asp, Glu, Cys, other amino acids). Circles show data for individual peaks, straight lines show model fits, and shaded regions show standard errors based on a linear model (Spearman's rank correlation for Asp peaks $\rho$=0.47, $p$=0.027, $N$=40; for Glu peaks $\rho$= 0.67, $p=2.15\times10^{-5}$, $N$=34; for Cys peaks $\rho$ = 0.92, $p<10^{-16}$, $N$=43; for other peaks $\rho$= 0.31, $p=1.56\times10^{-10}$, $N$=397).

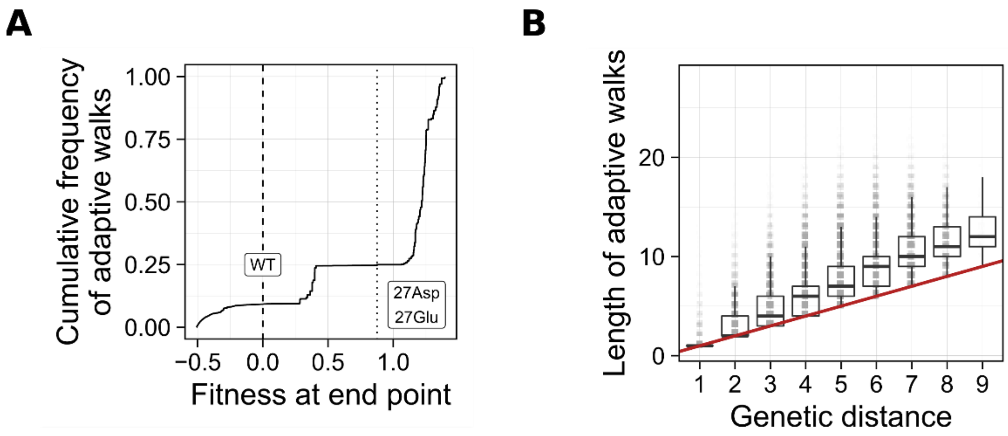


**Fig. S13. Adaptive walks with equal fixation probability for all fitness-increasing mutations.** The panels show the results for $10^6$ adaptive walks starting from randomly selected variants in the landscape. In contrast to the simulations presented on Fig 4, E and F, for which we used Kimura's fixation probability (favoring high fitness mutants), here we set fixation probability equal for all fitness-increasing mutations. **(A) Most evolving populations reach high fitness peaks.** The panel shows the cumulative distribution of fitness values at the end of adaptive walks ($N$=$10^6$ walks). The dashed vertical line $x$=0 shows the fitness value of the wild type. The dotted vertical line $x$=0.87 shows the fitness of the lowest among all high fitness peaks (27Asp/Glu). 75% of adaptive walks reached fitness 0.87 or higher. **(B) Adaptive walks reaching high fitness peaks are short.** The vertical axis shows the length of adaptive walks (number of mutations) that started from a randomly selected variant and reached a high fitness peak. The horizontal axis shows the (shortest) genetic distances between each starting variant and the attained peak. The red line $y$=$x$ shows the length of the shortest possible path to a high fitness peak, which is given by the genetic distance. Each box spans the interquartile range (IQR), each horizontal line inside a box indicates the median value, and each whisker extends to the minimum/maximum value outside the box within a 1.5 IQR interval.

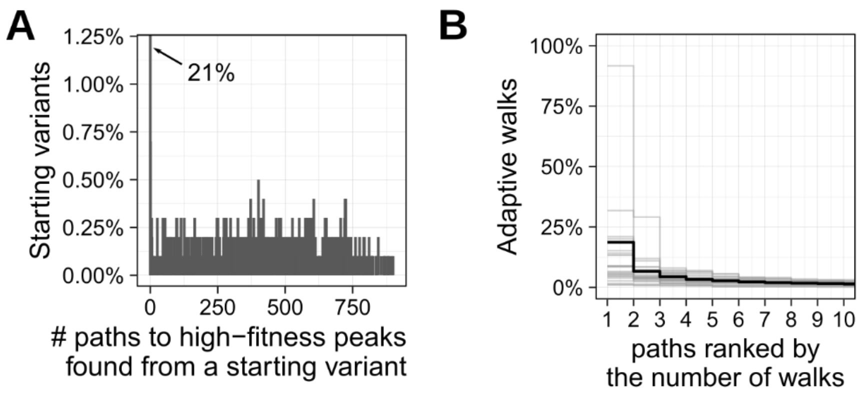


**Fig. S14. Evolutionary contingency. (A) Adaptive walks starting from the same variant follow different paths.** Distribution of the number of unique paths taken by a total of 1000 adaptive walks that start from the same variant. For this analysis, we used 1000 starting variants, i.e., we performed a total of $10^6$ adaptive walks. Walks initiated from 21% (209/1000) variants found no paths to any high fitness peaks, as indicated by the bar at $x$=0. For the remaining starting variants, adaptive walks followed multiple high fitness peaks. **(B) Adaptive walks preferably used some paths.** The vertical axis shows the percentage of adaptive walks (out of 1000), which take the most frequently used paths (horizontal axis). Each grey line summarizes the results for 1000 adaptive walks started from the same variant. To reduce visual clutter, grey lines are shown for only 24 randomly selected starting variants (24 grey lines) from which multiple high fitness peaks are reached. The black bold line shows the mean value for all 1000 starting variants.

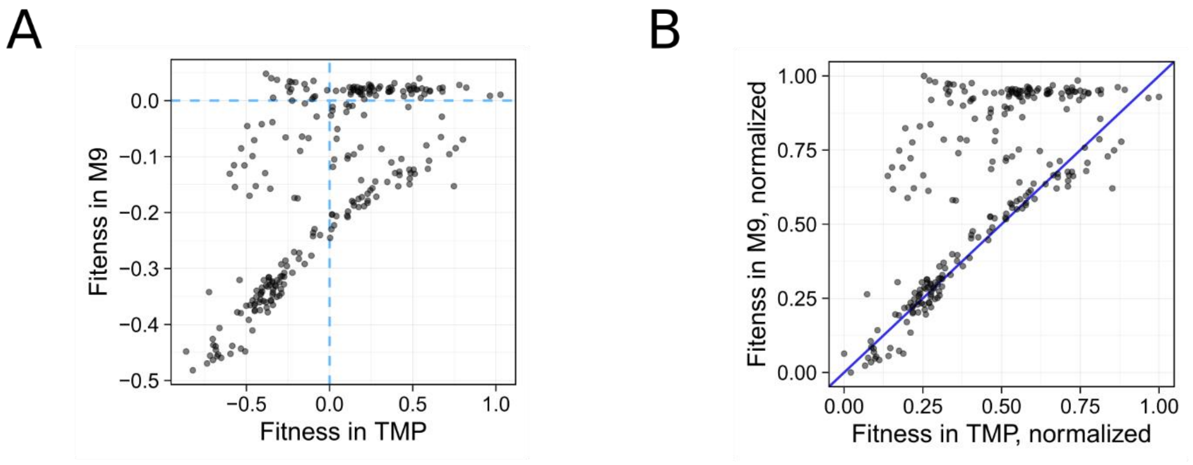


**Fig. S15 Fitness of DHFR variants in the presence and absence of trimethoprim. (A)** The panel shows the fitness of 254 DHFR variants in the presence of trimethoprim (0.4 mg/ml) on the horizontal axis, and the fitness of the same variants in M9 medium without trimethoprim on the vertical axis. The blue dashed lines at $x$=0 and $y$=0 show the fitness of the wildtype DHFR. **(B)** The same data as in panel A, but after scaling fitness values between 0 and 1. The blue line $x$=$y$ indicates the scenario where the relative fitness in the presence of trimethoprim matches that in the absence of trimethoprim. Note that most DHFR variants are above this line, indicating lower fitness in trimethoprim than expected based on their growth in the antibiotic-free environment. Only a few variants are below the line, indicating a trade-off in which a fitness advantage in the presence of antibiotics entails a fitness cost in the absence of antibiotic.

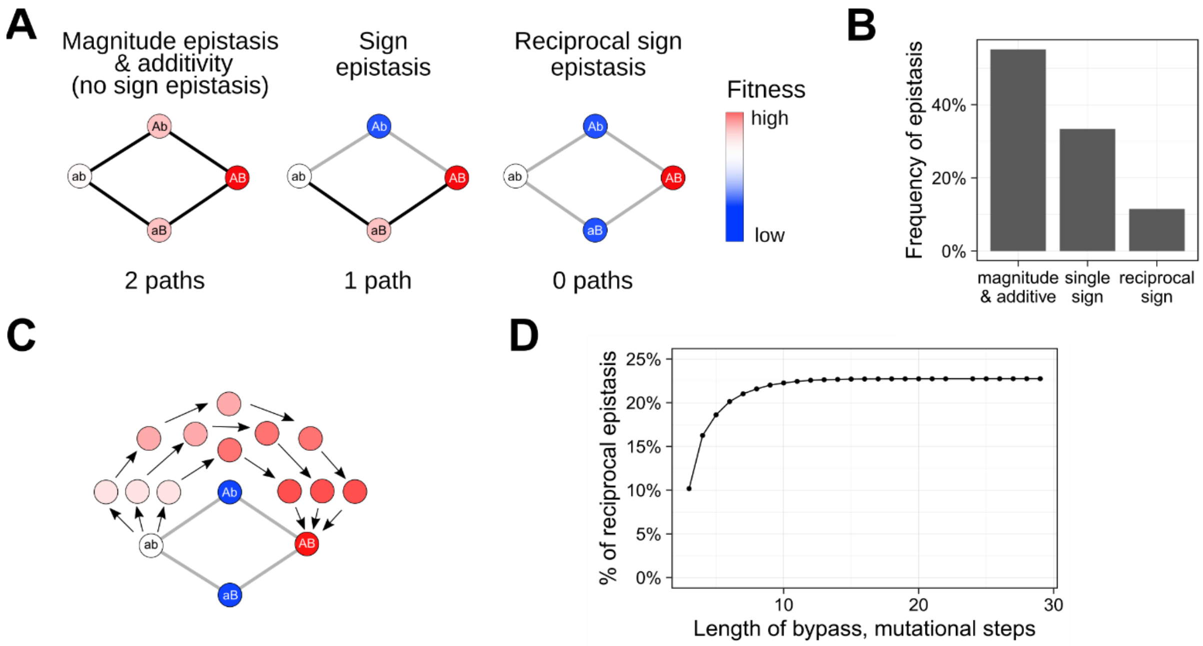


**Fig. S16. Epistasis. (A) Graph motifs representing three types of epistasis.** Each motif consists of four variants (circles), including a pair of genotypes *ab* and *AB* differing at two nucleotides (genetic distance: 2) and two intermediate genotypes *Ab* and *aB*. The color of circles indicates fitness (red = high fitness, white = intermediate fitness, blue = low fitness). Lines connecting genotypes show evolutionary paths. The black lines show accessible paths from variant *aa* to variant *AA*, and the grey lines show inaccessible paths, which are blocked by low fitness of intermediate variants (*Ab* and/or *aB*). The three principal type of epistasis shown here differ in the number of accessible paths between variant *ab* and *AB*. **(B) Prevalence of three types of epistasis.** The barplot shows the prevalence of three types of motives in our landscape, as explained from panel A (*N*=740,211 double mutant pairs). **(C) Schematic illustration of the principle behind an extradimensional bypass.** Four genotypes (ab, Ab, aB, AB) connected by grey lines indicate a hypothetical case of reciprocal sign epistasis. No direct accessible paths exist between variants *ab* and *AB* due to the low fitness of the intermediate variants *Ab* and *aB* (blue circles). However, variant *AB* may be accessible from *ab* via indirect paths. Indirect accessible paths are longer than direct paths, because they require mutation of additional nucleotides or sites, but the fitness of variants along these paths increases monotonically from *ab* (white circle) to *AB* (dark red circle), which makes these paths evolutionary accessible (black arrows). **(D) Extradimensional bypasses exist for only a minority of cases of reciprocal sign epistasis.** The horizontal axis shows the maximal length of an indirect path (extradimensional bypass), and the vertical axis shows the cumulative percentage of double mutant pairs with reciprocal sign epistasis for which a bypass of that maximal length exists in the landscape. The total number of squares with reciprocal sign epistasis is 85,203.

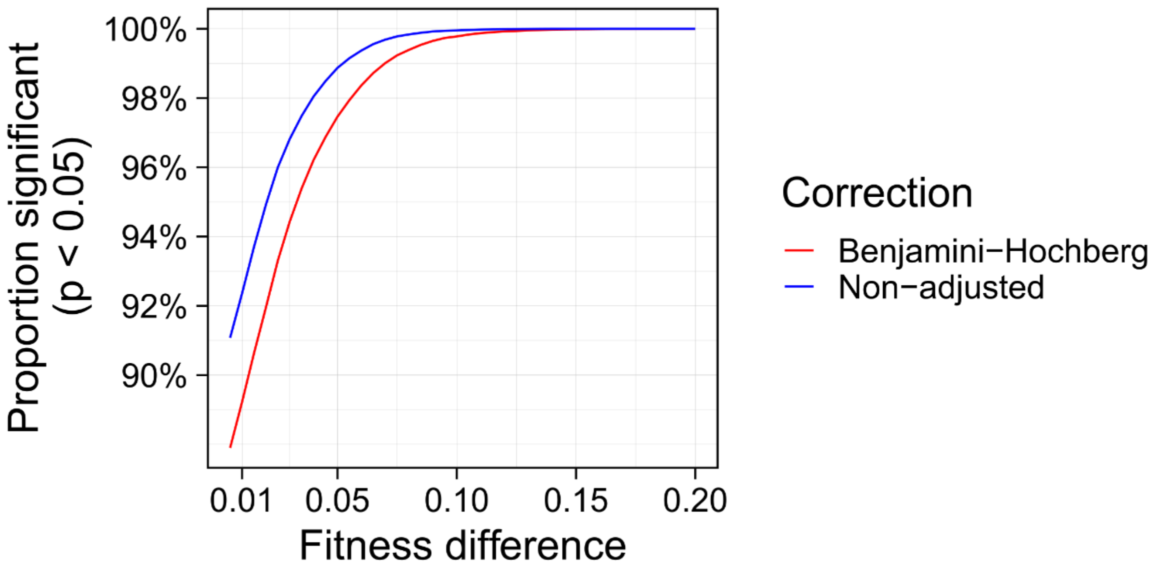


**Fig. S17. Significance of fitness effect in the Wald test.** The proportion of statistically significant fitness differences between connected variants in the landscape. The blue line shows unadjusted p-values obtained using the Wald test after fitting a generalized linear model. The red line shows the adjusted p-values using a Benjamini-Hochberg correction ($N$=45,731,140, $N$ is the total number of tests using all pairs of variants at a genetic distance not exceeding two nucleotide substitutions).

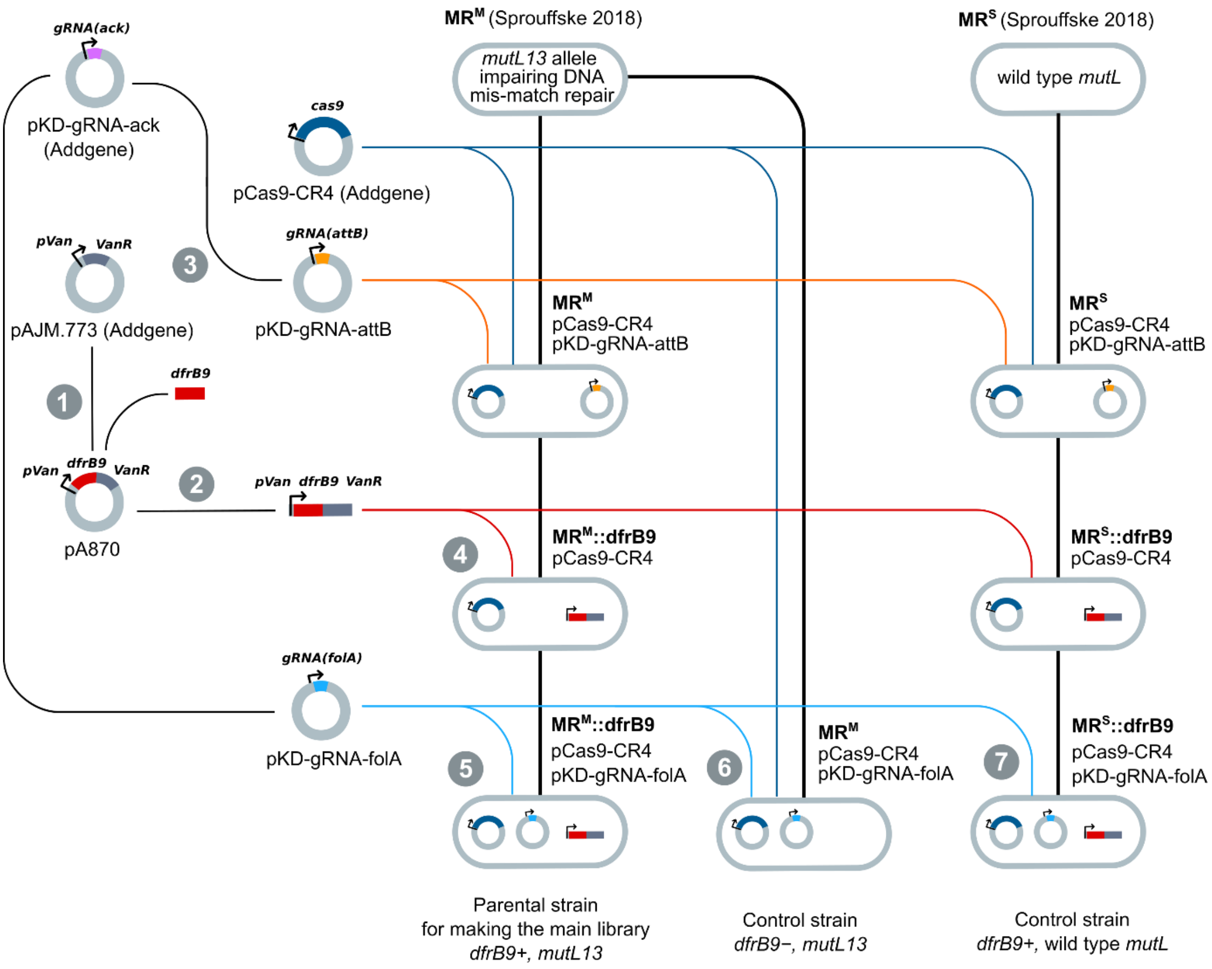


**Fig. S18. Construction of parental strains.** The diagram shows the creation of one parental strain (MR$^{M}$::*dfrB9*/pCas9-CR4/pKD-gRNA-*folA*) used in the main experiment and two control strains (MR$^{M}$/pCas9-CR4/pKD-gRNA-*folA* and MR$^{S}$::*dfrB9*/pCas9-CR4/pKD-gRNA-*folA*, bottom row). We derived these strains from the two strains MR$^{M}$ and MR$^{S}$ from a previous study[80] (top row). MR$^{M}$ has *mutL13* allele impairing DNA mismatch repair and MR$^{S}$ has the wild type *mutL* gene. For construction, we obtained three vectors from Addgene (pCas9-CR4, pKDsgRNA-ack, pAJM.773), and constructed three other vectors as detailed in Methods. Numbers in gray circles correspond to the following sections in Methods, which explain each construction step: (1) Cloning *dfrB9* under the control of the VanR regulator, (2) Generating a template for chromosomal integration of *dfrB9*, (3) Construction of pKD-gRNA-*attB* to edit the *attB* site, (4) Chromosomal integration of *dfrB9* using gene editing, (5) Constructing the parental strain, (6) Creating control parental strain without *dfrB9* integration, (7) Creating parental strain with the wild type *mutL*.

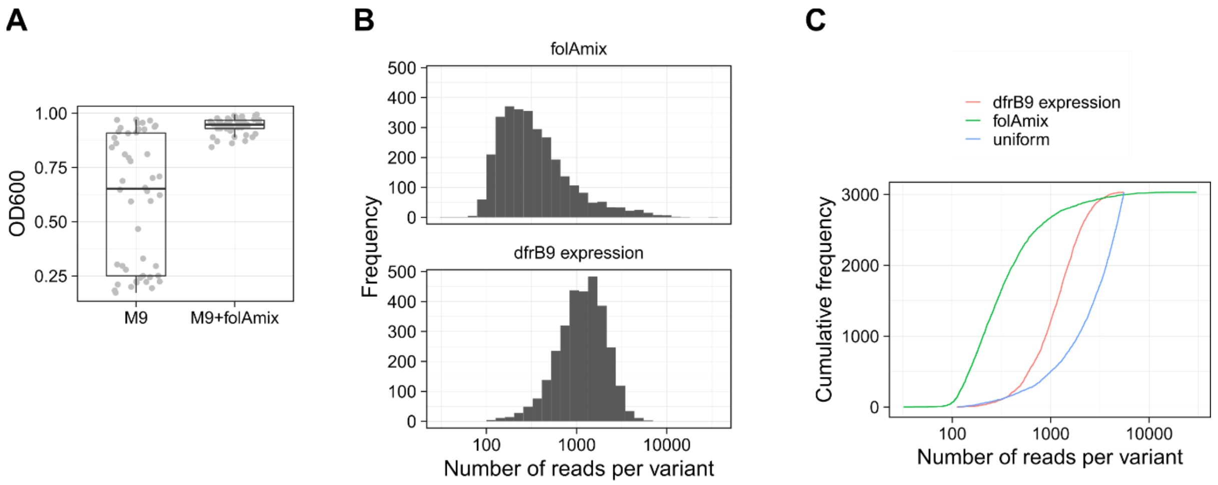


**Fig. S19. (A)** The effect of folA-mix on the growth of *folA* mutants (*N*=48 mutants)**. (B)** The comparison of variant frequencies in mutant libraries obtained using two different strategies for compensating for a lack of *folA* function in low fitness mutants. The top panel shows the frequency of variants in the library obtained by performing Cas9 mutagenesis in the presence of folA-mix. The bottom panel shows the frequency of variants in the main library, in which the *dfrB9* gene was expressed. For comparison, the data from the main library (bottom panel) was subsampled to contain an equal number of variants as the library obtained using folA-mix (*N*=3031). **(C)** Cumulative distributions of variant frequencies in both libraries. The green curve shows this distribution for the library obtained using folA-mix (*N*=3031 variants), the red curve shows the library obtained by expressing the *dfrB9* gene (subsampled to comprise *N*=3031 variants), and the blue line shows a simulated distribution obtained under the assumption that all variants have equal frequencies.

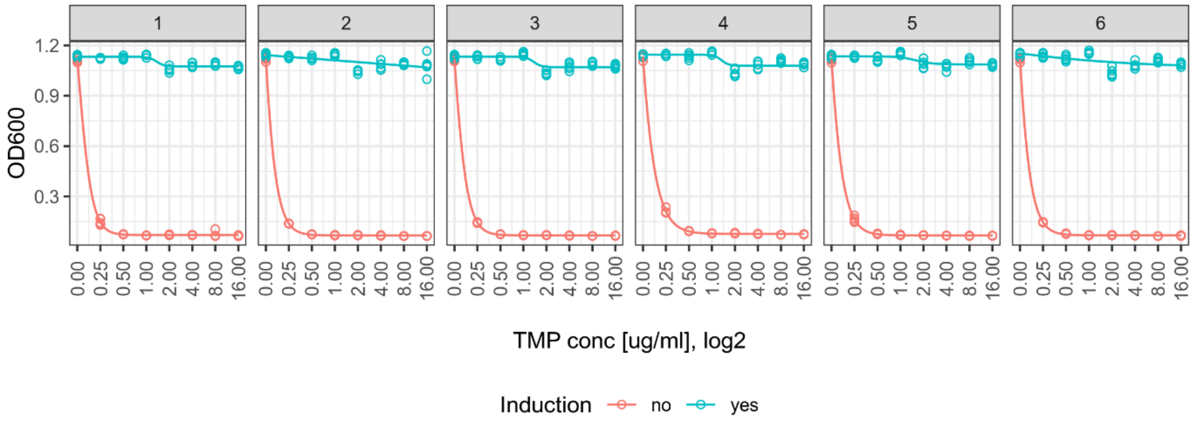


**Fig. S20. Trimethoprim resistance of strain MR[M] carrying the gene *dfrB9* on plasmid pA870.** The vertical axis indicates bacterial growth (optical density at 600 nm, $OD_{600}$) after 24 h, for different trimethoprim concentrations (horizontal axes). Turquoise circles show data from cultures grown in the presence of the inducer of *dfrB9* expression (10 µM vanillic acid), and red circles correspond to cultures grown in the absence of the inducer. We performed the experiment for six independently isolated clones harboring the *dfrB9* gene, as indicated in the labels above each of the six panels. We used six replicate cultures, each shown as an individual circle, per combination of clone, presence/absence of inducer, and trimethoprim concentration.

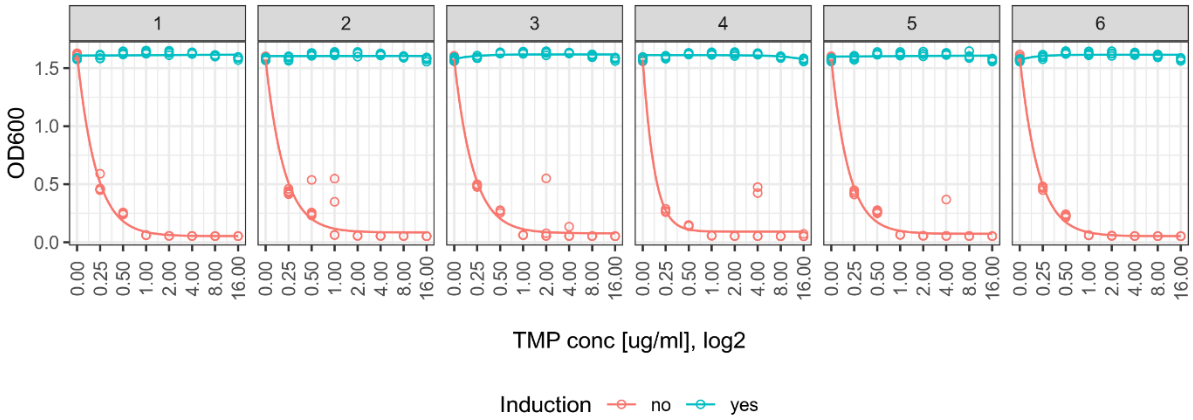


**Fig. S21. Trimethoprim resistance of strain MR$^{M}$ containing the gene *dfrB9* integrated into the chromosome.** The vertical axis shows the optical density of bacterial cultures at $\lambda$=600 nm ($OD_{600}$) measured after 24 h of growth at the different trimethoprim concentrations indicated on the horizontal axis. Turquoise circles show data from cultures grown in the presence of the inducer of *dfrB9* expression (10 µM vanillic acid), and red circles correspond to cultures grown in the absence of the inducer. We performed the experiment for six independently isolated clones with the integrated *dfrB9* gene, as indicated in the labels above each of the six panels. We used six replicate cultures, each shown as an individual circle, per clone, presence/absence of the inducer, and trimethoprim concentration combination.

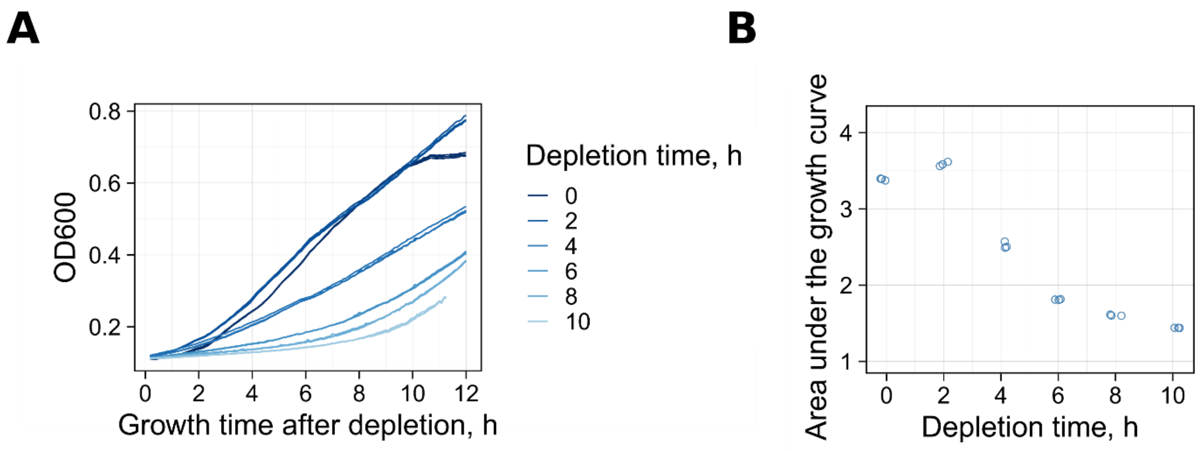


**Fig. S22. Effect of DfrB9 depletion on library growth. (A)** The growth of the mutant library after DfrB9 depletion. We depleted cultures by growing them without adding vanillic acid, the inducer of *dfrB9* expression. The vertical axis shows the optical density at $\lambda$=600 nm ($OD_{600}$) of DfrB9-depletion depleted cultures transferred to the fresh medium M9 with 0.4 µg/ml trimethoprim. We measured optical density at 5-minute intervals. The line color indicates different durations of depletion (3 replicate curves for each duration). **(B)** Summary of the growth data from panel **A** as a function of depletion time (horizontal axis). The vertical axis shows the area under the growth curve calculated from the data in panel **A**. Library growth decreases substantially after a few hours of DfrB9 depletion, because most variants have non-functional or low activity DHFR.

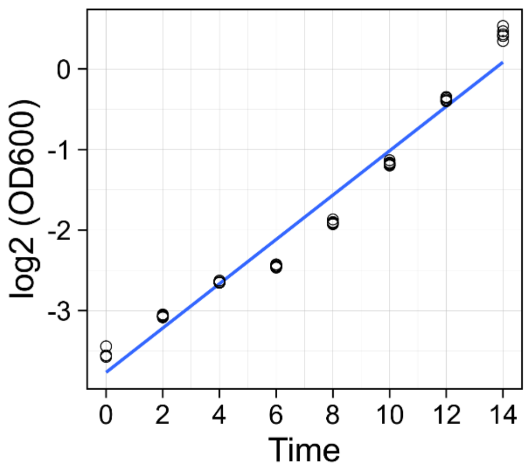


**Fig. S23. The growth curve from the mass selection experiments.** We measured optical density at $\lambda$=600 nm ($OD_{600}$) at 2 h intervals during mass selection experiments in 6 replicate cultures (individual circles). Note the $log_2$-transformed vertical axis. The systematic deviation from a linear regression line (blue) reveals a modest non-linearity in the growth dynamics.

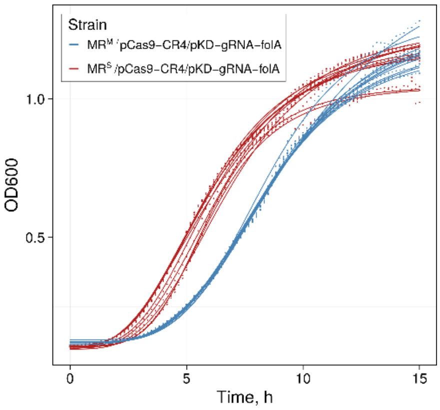


**Fig. S24. Growth of MR$^{M}$ strain (with mutL13) and MR$^{S}$ (wildtype *mutL*).** The vertical axis shows the optical densities ($\lambda$=600 nm) of bacterial cultures measured at a ten-minute interval for 15 hours. Circles show individual data points from 10 replicate cultures. Blue curves indicate data from the MR$^{M}$ strain with the mutL13 allele which impairs DNA mismatch repair. Red curves indicate data from the isogenic strain MR$^{S}$, which carries the wild-type version of the *mutL* gene.

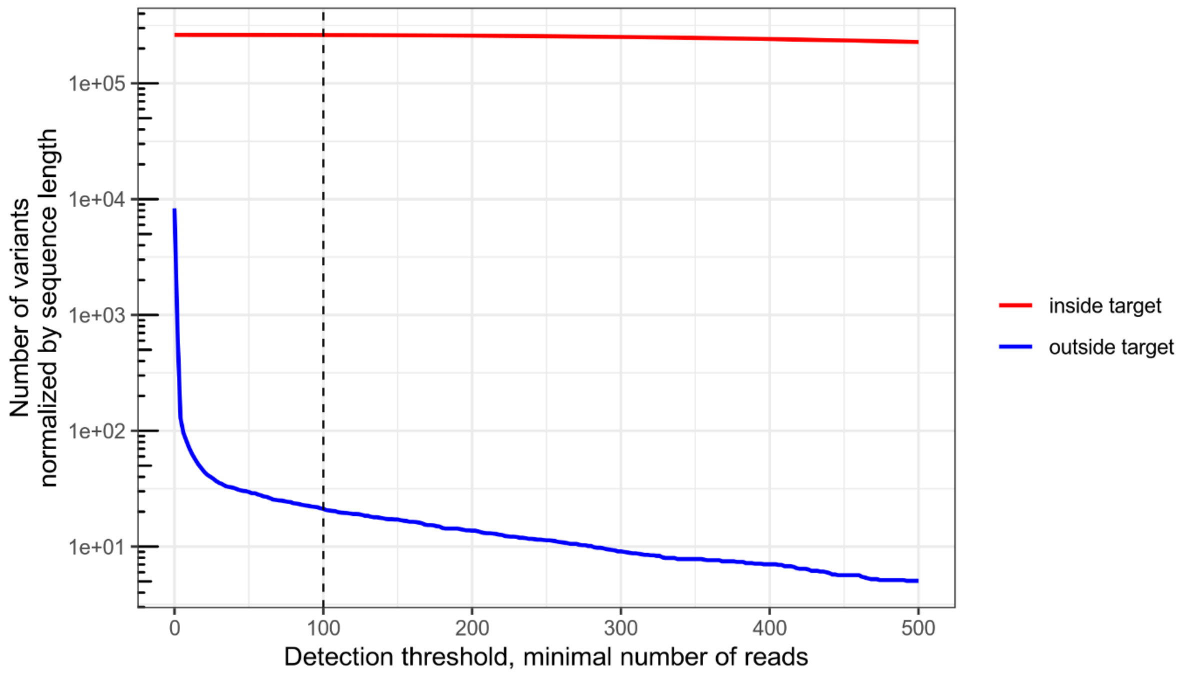


**Fig. S25 False positive rate for detecting DNA sequence variants.** We determined the number of unique DHFR variants in our library by setting a minimal number of reads as a detection threshold. The blue line shows variants detected outside of the mutagenized region. Likely, such variants are sequencing errors (false positives, i.e., reads that are incorrectly assigned to a variant due to sequencing errors). The red line shows the number of variants detected inside of mutagenized regions including both true positives and false positives. This means that the total read counts for any variant contains a certain fraction of false positive reads, which may potentially bias the estimate of true read counts (i.e., reads correctly assigned to a variant). However, the comparison of blue and red lines suggests that at high coverage, false positive reads represent only a very small fraction of total counts, resulting in a small bias. We chose the conservative threshold of 100 reads, at which the expected fraction of false positive reads is just 22/261382 = 0.000085.

### Supplementary Tables

**Table S1**. Vectors used in this study

| **Name** | **Genotype** | **Selective antibiotics (concentration µg/ml)** | **Source** | **T, °C** | **Description** |
|---|---|---|---|---|---|
| pKDsgRNA-ack | exo, beta, gam, sgRNA-ack | Spectinomycin (50) | Addgene, 62654 [44,81] | 30 | Vector for expressing gRNA targeting ack |
| pCas9-CR4 | cas9-CR4 | Chloramphenicol (34) | Addgene, 62655 [44,81] | 37 | Vector for expressing cas9 |
| pKD-gRNA-folA | exo, beta, gam, sgRNA(folA) | Spectinomycin (50) | This study | 30 | Vector for expressing gRNA targeting *folA* |
| pKD-gRNA-attB | exo, beta, gam, sgRNA(attB) | Spectinomycin (50) | This study | 30 | Vector for expressing gRNA targeting attB |
| pAJM.773 | vanR | Kanamycin (50) | Addgene 108527[82] | 37 | Vector for cloning *dfrB9* gene |
| pA870 | dfrB, vanR | Kanamycin (50) | This study | 37 | pAJM.773 carrying *dfrB9* |

**Table S2**. *E. coli* strains used in this study

| **Name** | **Genotype** | **Selective antibiotics (concentration µg/ml)** | **Source** | **T, °C** | **Description** |
|---|---|---|---|---|---|
| **E. coli strains** | | | | | |
| $MR^M$ | F-, fhuA2, lacY1, tsx-1 or tsx-70, glnV44 (AS), gal-6, λ-, xyl-7, mtlA2, mutL13 | | Yale's Coli Genetic Stock #4816 [80,83,84] | 37 | Derivative of K-12 MG1655 with impaired mismatch repair (*mutL13*). Amber (UAG) suppressor |
| $MR^S$ | F-, fhuA2, lacY1, tsx-1 or tsx-70, glnV44 (AS), gal-6, λ-, xyl-7, mtlA2, cycA30::Tn10 | | Wagner's lab stock[80] | 37 | Derivative of K-12 MG1655 with wild type *mutL*. Amber (UAG) suppressor |
| $MR^M$ /pCas9-CR4/ pKD-gRNA-attB | cas9-CR4, exo, beta, gam, sgRNA-attB | Spectinomycin (50) Chloramphenicol (34) | This study | 30 | $MR^M$ strain ready for chromosomal integration at attB locus |
| $MR^M$ /pCas9-CR4/ pKD-gRNA-folA | cas9-CR4, exo, beta, gam, sgRNA-folA | Spectinomycin (50) Chloramphenicol (34) | This study | 30 | Control parental strain for creating mutant libraries without *dfrB9* |
| $MR^M$::dfrB9/pCas9-CR4 | cas9-CR4, *dfrB9* | Chloramphenicol (34) | This study | 37 | $MR^M$ with integrated *dfrB9* gene |
| $MR^M$::dfrB9/pCas9-CR4/ pKD-gRNA-*folA* | cas9-CR4, exo, beta, gam, sgRNA-folA, *dfrB9* | Spectinomycin (50) Chloramphenicol (34) | This study | 30 | Parental strain for creating |

| | | | | | |
|---|---|---|---|---|---|
| | | | | | mutant libraries |
| MR$^S$::dfrB9/pCas9-CR4/ pKD-gRNA-*folA* | cas9-CR4, exo, beta, gam, sgRNA-folA, *dfrB9* | Spectinomycin (50) Chloramphenicol (34) | This study | 30 | Control parental strain for creating mutant libraries MG1655 with the wild type *mutL* |
| DH5-α | | | Escudero's lab stock | 37 | Strain for molecular cloning |
| DH5-α-pA870 | | Kanamycin (50) | This study | | DH5-α carrying pA870 |

**Table S3**. Oligonucleotides used in this study. Capital letters in the second column show degenerate nucleotides (N = A, T, G, C and R = A, G).

| Name | Sequence |
|---|---|
| dfrB9-pVan-For | agaaagaggggaaatactagttaggccccgcaatgcgaac |
| dfrB9-pVan-Rev | ttctggaatttgcgctgtctagagcctaacgttcgaggtg |
| bb-Marionette-F | caaattccagaaaagaggcc |
| bb-Marionette-R | ctagtatttcccctctttctc |
| pr009 | tttataacctccttagagctcga |
| pr010 | ccaattgtccatattgcatca |
| pr014 | attgtaatgcggcgagtcca |
| pr015 | caccgacttcacccacgtta |
| pr049 | ttaaaccaggcgagatcggcgttttagagctagaaatagcaag |
| pr050 | gccgatctcgcctggtttaagtgctcagtatctctatcactga |
| pr068 | tcaagttagtataaaaaagcgttttagagctagaaatagcaag |
| pr069 | gcttttttatactaacttgagtgctcagtatctctatcactga |
| pBAD-rev | gatttaatctgtatcagg |
| pr076 | gttcacaggttgctccgggctatgaaatagaaaaatgaatccgttgaagctggtctcccgcttaacgat |
| pr077 | gtattaaaaacaactttttgtctttttaccttcccgtttcgctcaagttagtaacgaagcagggattctgcaa |
| pr078 | cctggtgtcagggatgcaaa |
| pr082 | gccgaaaggcgaaactcatc |
| ngs_folA_F01 | agtcattgagtcagtctgattgcggcgtta |
| ngs_folA_F02 | gatctcattctcagtctgattgcggcgtta |
| ngs_folA_F03 | cgcttatccttcagtctgattgcggcgtta |
| ngs_folA_F04 | tatcatgcagtcagtctgattgcggcgtta |
| ngs_folA_F05 | atctgcgtactcagtctgattgcggcgtta |
| ngs_folA_F06 | gattgcacgctcagtctgattgcggcgtta |
| ngs_folA_R | ggacgaccgattgattccca |
| t022 | ccataatcacgggtttatttaaggtgttgcgtttaaaccaggcNNNNNNNNNaggcaggttccacggcatggcgttttccatg |
| t023 | ccataatcacgggtttatttaaggtgttgcgtttaaaccaggcNNNNtcggcaggcaggttccacggcatggcgttttccatg |

**Table S4**. Sequence analysis pipeline statistics

| Pipeline step | Number of reads | Fraction retained from raw data | Fraction retained from previous step |
|---|---|---|---|
| ***Before selection*** | | | |
| 1. Raw data | 847212668 | 1.0000000 | 1.0000000 |
| 2. De-multiplexing | 817697668 | 0.9651622 | 0.9651622 |
| 3. Trimming | 748968878 | 0.8840388 | 0.9159484 |
| 4. Merging | 374429821 | 0.8839099 | 0.9998542 |
| 5. Aligning | 374426014 | 0.8839009 | 0.9999898 |
| 6. De-noising | 345070984 | 0.8146030 | 0.9215999 |
| ***After selection*** | | | |
| 1. Raw data | 457433858 | 1.0000000 | 1.0000000 |
| 2. Demultiplexing | 448725376 | 0.9809623 | 0.9809623 |
| 3. Trimming | 428403058 | 0.9365355 | 0.9547110 |
| 4. Merging | 214196772 | 0.9365147 | 0.9999778 |
| 5. Aligning | 214196545 | 0.9365137 | 0.9999989 |
| 6. Denoising | 208879397 | 0.9132660 | 0.9751763 |

### Methods

#### Strains and plasmids

*E. coli* strains $MR^M$ and $MR^S$ are derivatives of K-12 MG1655[80]. Strain $MR^M$ ('M' for medium mutation rate) has an increased mutation rate due to the mutation *mutL13* in the gene encoding the DNA mismatch repair protein MutL[80]. The mutation rate of $MR^M$ is 16 times higher than that of the isogenic strain $MR^S$ ('S' for small mutation rate) in which the *mutL* gene has the wild type genotype[80]. Both strains also carry the amber stop codon suppression mutation glnV44. We obtained $MR^M$ and $MR^S$ strains from Wagner laboratory stocks.

We used the plasmid vectors pKDsgRNA-ack and pCas9-CR4 for gene editing[44]. We used the plasmid pAJM.773 to clone the *dfrB9* gene[82]. We obtained these plasmids from the Addgene plasmid repository. All vectors and strains used in this study are listed in Tables S1 and S2 and shown in Fig. S18.

#### Media and reagents

To prepare SOB medium we dissolved 25.5g of its solid stock (VWR J906) in 960 ml of water, and autoclaved the medium before use. To prepare SOC medium, we added 20 ml of 1 M D-glucose (Sigma G8270), and 20 ml of 1 M magnesium sulfate (Sigma 230391) to 960 ml of SOB medium.

We purchased M9 minimal salt from Sigma (M6030), dissolved it according to the supplier's instructions, sterilized the solution by autoclaving, and supplemented it with 0.4% glucose (Sigma G8270), 0.2% casamino acid (Merk Millipore, 2240), 2 mM magnesium sulfate (Sigma 230391), and 0.1 mM calcium chloride (Sigma C7902).

To prepare M9 folA-mix or SOB folA-mix, we supplemented SOB and M9 with 0.002 mM calcium-D-pantothenate (PHR1232), 0.5mM glycine (G6761), 0.5 mM L-methionine (M5308), 1mM thymidine (T9250) and 1.25 mM adenine (A9126). We purchased all the supplements from Sigma, dissolved them in water and filter-sterilized before use.

Where necessary, we supplemented growth media with 0.4 µg/ml trimethoprim (BioVision B1539), 50 µg/ml spectinomycin (Sigma S4014), 34 µg/ml chloramphenicol (Sigma C1919), 50 µg/ml kanamycin (Sigma BP861), as well as with 0.2% L-arabinose (Sigma A3256), 0.16 µg/ml anhydrotetracycline (Cyaman 10009542), and/or 10 µM vanillic acid (Sigma 94770).

#### Preparation of electro-competent cells using the mannitol-glycerol method

We prepared electro-competent cells using a protocol adapted from Warren[44,85]. Specifically, we diluted an overnight culture 100-fold in 4 ml of SOB medium, and incubated the culture until the cell density reached $OD_{600}$ = 0.5. We chilled the cells on ice for 6 min and pelleted them by centrifuging (10 min at 3000 × g, 4°C). After removing the supernatant, we resuspended the cells in 1 ml of ice-cold water and transferred the suspension to a 2 ml microcentrifuge tube. We then gently added 1 ml of glycerol-mannitol solution (20% glycerol, 1.5% mannitol) to the bottom of the tube. We centrifuged the tubes for 10 min at 3000 g, 4°C, and carefully removed the supernatant. Subsequently, we resuspended the cell pellet in 100-200 µl of glycerol-mannitol solution and used freshly prepared competent cells for electroporation.

### Design of guide RNA (gRNA) spacers

We used the R package `Crisprseek`[86] to find sequences of potential gRNA spacers for editing the *E. coli* genome (GenBank accession number NC_000913). We used the sequences of identified candidate gRNA spacers to predict Cas9 activity with the help of a program published by Guo et al[87]. We selected the candidate gRNA with the highest predicted activity for gene editing.

### Chromosomal integration of *dfrB9* using gene editing

<u>*The rationale for integrating dfrB9.*</u> DHFR is essential for *E. coli* growth in minimal medium and shows low tolerance to mutations[48,76]. This makes it challenging to maintain low activity or nonfunctional DHFR variants in a population, which is required to study a DHFR library that is combinatorially complete. Previous studies used the so called folA-mix, a medium complementing the metabolic deficiencies of *folA* mutants for this purpose[76,88–91]. Although this medium improves the growth of mutants (Fig. S19A), it does not fully rescue their fitness, resulting in a highly skewed representation of variants after mutagenesis in the presence of folA-mix (see Fig. S19, B and C, section *The effect of folA-mix on the uniformity of variant frequencies after gene editing*). The skewed representation dramatically reduces the ability to estimate variant fitness for many variants, because most sequencing reads would represent a few abundant variants, while low frequency variants would have low coverage.

To allow the persistence of DHFR variants with low or no functionality in a population, we constructed an *E.coli* strain carrying *dfrB9*, an alternative gene encoding dihydrofolate reductase (GenBank accession number NG_052167). *dfrB9* is a member of the DfrB family of dihydrofolate reductases, which have an independent evolutionary origin from the *E.coli* chromosomal *folA* gene[92,93]. DfrB9 can replace the function of chromosomally encoded dihydrofolate reductase and is intrinsically resistant to the antibiotic trimethoprim. We integrated the *dfrB9* gene into the chromosome of the $MR^M$ strain following several steps described in the next sections.

<u>*Cloning dfrB9 under the control of the VanR regulator.*</u> We cloned the *dfrB9* gene into the pAJM.773 plasmid vector under the control of the synthetic VanR regulator, which activates expression in the presence of vanillic acid[82]. The *dfrB9* gene cassette was synthesized using a commercial service (IDT, USA). We amplified the cassette with the oligonucleotides dfrB9-pVan-For and dfrB9-pVan-Rev (Table S3) using a polymerase chain reaction (PCR) with the following cycling protocol: 98°C for 3 min, 36 cycles of 98°C for 10 s, 55°C for 20 s, and 72°C for 1 min 30 s, followed by 5 min at 72°C. The PCR reaction contained Phusion HF buffer, 0.02 U/µl Phusion DNA polymerase (NEB M0530), 0.2 mM nucleotides, and 0.5 µM primers. We amplified the plasmid backbone with the primers bb-Marionette-F and bb-Marionette-R (Table S3). To clone the amplified products, we performed Gibson assembly[94] using Gibson Assembly Master Mix (NEB E2611). We transformed the assembly reaction by electroporation into *E. coli* DH5-α (1 mm cuvette, 2.4 kV, BioRad MicroPulser), and selected for successful transformants on SOC agar plates containing 50 µg/ml kanamycin. We called the resulting plasmid p870 and verified its correct sequence by Sanger sequencing (Microsynth AG, Switzerland).

We verified the expression of *dfrB9* from the p870 vector in six independent clones harboring the vector. Specifically, we diluted overnight cultures of these clones 1000-fold and inoculated each dilution into M9 medium containing a range of trimethoprim concentrations (0, 0.25, 0.5, 1, 2, 4, 8, 16 µg/ml). We included two treatments, one with the inducer (10 µM vanillic acid) and one without. Each treatment had six replicate cultures per clone and trimethoprim concentration. We grew cultures in 96-well plates (TPP 92096) at 30°C with shaking at 1000 rpm in a Stuart SI505 incubator (Cole-Parmer, UK). After 24 h, we measured the optical density ($\lambda$=600 nm) of the cultures using a plate reader (Spark, Tecan). As expected, we found a high level of resistance to trimethoprim in the induced cultures due to the expression of DfrB9 (see Fig. S20).

Generating a template for chromosomal integration of *dfrB9*. To generate a recombination template for chromosomal integration of *dfrB9*, we used a PCR. Specifically, we amplified a 1.6 kb fragment from the p870 vector, which included *dfrB9*, as well as the pVan promoter upstream and the downstream regulator gene *vanR*[82,95]. The forward and reverse primers for this amplification (pr076 and pr077, Table S3) contained overhangs homologous to the sequences flanking the *attB* site in the *E. coli* MG1655 genome. We performed the PCR amplification with the following program: 98°C for 3 min, 36 cycles of 98°C for 10 s, 55°C for 20s, and 72°C for 1 min 30 s, followed by 72°C for 5 min. The reaction contained Phusion HF buffer, 0.02 U/µl Phusion DNA polymerase (NEB M0530), 0.2 mM nucleotides, and 0.5 µM primers. We purified the PCR product from a 1.2% agarose gel using the QIAquick PCR & Gel Cleanup Kit (Qiagen 28506). The purified product served as a recombination template for homologous repair.

Construction of pKDsgRNA-attB to edit the attB site. As a site for inserting *dfrB9* into the chromosome, we chose the attB site[96], the locus for site-specific integration of the bacteriophage lambda, which is commonly used for lambda-based integration vectors[97,98]. To express a guide RNA that targets this site, we constructed the pKDsgRNA-attB vector by replacing the spacer region of the gRNA gene in pKDsgRNA-ack using circular polymerase extension cloning[44,99]. To this end, we first amplified two overlapping fragments of pKDsgRNA-ack, using primer pairs pr009/pr068 and pr010/pr069 (Table S3). To replace the spacer, the overhangs in the primers pr068 and pr069 contain 20 nucleotide (nt) sequences homologous to the attB site of the chromosome (GenBank NC_000913.3, positions 807328-807342). We performed the PCR reactions using 0.02 U/µl Phusion DNA polymerase in Phusion HF buffer (NEB M0530), with 0.5 µM of each primer and 0.2 mM nucleotides. The program consisted of denaturation 98°C for 3 min, 36 cycles of 98°C for 10 s, 55°C for 20 s, and 72°C for 1 min 30s, followed by extension at 72°C for 5 min. We treated the amplified fragments with 20 units DpnI (NEB R0176) and 20 units ExoI (NEB M0293S) for 1 h at 37°C and purified them using the Monarch PCR & DNA Cleanup Kit (NEB T1030S). We then combined the two purified fragments using a single circular polymerase extension reaction[99]. The reaction contained Phusion HF buffer, 0.02 U/µl Phusion DNA polymerase (NEB M0530), 0.2 mM nucleotides, and 0.08 pM of DNA fragments. We used the following program for the circular polymerase extension reaction: 98°C for 30 s, 15 cycles of 98°C for10 s, 55°C for 30 s, and 72°C for 2 min, followed by 72°C for 5 min. Subsequently, we transformed the strain $MR^M$ carrying pCas9-CR4 with 2 µl of the reaction product using electroporation (1 mm cuvette, 2.4 kV, BioRad MicroPulser). To recover the transformed cells, we incubated them in SOC medium for 2 h at 30°C, and plated them on SOC agar plates with 50 µg/ml spectinomycin and 34 µg/ml chloramphenicol. Selected colonies were expected to carry pKDsgRNA-attB. We picked and verified three candidate colonies by

Sanger sequencing (Microsynth AG, Switzerland) using the universal primer pBAD-rev (Table S3), and named the verified clones $MR^M$/pCas9-CR4/pKDsgRNA-attB.

*Verifying cas9 activity*. We used the same three clones of $MR^M$/pCas9-CR4/pKDsgRNA-attB to verify Cas9-induced mortality. We resuspended bacterial colonies in PBS buffer, diluted them $10^1$- $10^5$-fold, and plated this dilution series on SOB agar plates (50 µg/ml spectinomycin, 34 µg/ml chloramphenicol), with or without the inducer anhydrotetracycline (0.16 µg/ml). Anhydrotetracycline activates the expression of Cas9 and of the gRNA genes. Their gene products form the gRNA-Cas9 complex[100]. The gRNA-Cas9 complex binds to a targeted region (homologous to the spacer sequence of gRNA), and introduces a double strand break[101,102]. This double strand break is lethal unless a template is provided for homologous repair. We observed a $10^4$-fold reduction in colony forming units (CFU) counts on plates containing anhydrotetracycline compared to control plates, which indicated sufficient Cas9 activity[44].

*Chromosomal integration using gene editing.* We inserted the 1.6 kb double-stranded fragment amplified from pA870 (see section *Generating a template for chromosomal integration of dfrB9*) into the chromosome using the noSCAR protocol[44,81]. To this end, we prepared transformation competent cells ($MR^M$/pKDsgRNA-attB/pCas9-CR4) using the glycerol-mannitol method, and induced the lambda Red genes (expressed from pKDsgRNA-attB) with 0.2% arabinose for recombineering [103]. We then transformed the cells with 2 µg of the purified DNA fragment (1 mm cuvette, 2.4 kV, BioRad MicroPulser), and recovered them in SOC medium for two hours (30°C, 225 rpm). After recovery, we plated the cells on SOC agar plates (50 µg/ml spectinomycin, 34 µg/ml chloramphenicol, 0.16 µg/ml anhydrotetracycline), and incubated them at 37°C for 40 h. Cells expressing *dfrB9* should be resistant to trimethoprim. Therefore, to select cells with integrated *dfrB9*, we scraped the colonies from the agar plates, resuspended them in PBS, diluted $10^1$-$10^4$ times, and plated onto selective SOB agar plates containing 100 µM vanillic acid and 16 µg/ml trimethoprim. After 24 h incubation at 37°C, we picked six colonies to confirm integration.

*Confirming dfrB9 expression from the chromosome*. We confirmed *dfrB9* expression by growing candidate clones at high concentrations of trimethoprim (up to 16 µg/ml). Specifically, we prepared overnight cultures, and diluted them 1000-fold in M9 containing trimethoprim (0, 0.25, 0.5, 1, 2, 4, 8, 16 µg/ml), with or without the addition of 10 µM vanillic acid. For all experiments, we cultured six replicates for 24h at 30°C in 96-well plates (TPP 92096), using a Stuart SI505 incubator (Cole-Parmer, UK) at 1000 rpm. After incubation, we measured the optical density ($\lambda$=600 nm) using a Spark plate reader (Tecan). The induced cultures showed uninhibited growth across all trimethoprim concentrations. In contrast, in the absence of inducer, growth was completely inhibited by 4 µg/ml trimethoprim, with no evidence of leaky *dfrB9* expression (Fig. S21).

*Verification by sequencing.* We used genomic DNA isolated from candidate clones using the DNeasy Blood & Tissue Kits (Qiagen 69504) according to the manufacturer's protocol for gram-negative bacteria. This DNA served as a template to amplify the fragment spanning the entire region of *dfrB9* integration. The required PCR reaction was composed of 0.020 U/µl Phusion DNA polymerase in Phusion HF buffer (NEB M0530), 0.5 µM primers pr078 and pr082 (Table S3), as well as 0.2 mM nucleotides. We performed the PCR with the following thermocycling protocol: 98°C for 3 min, 36 cycles of 98°C for 10 s, 55°C for 30 s, 72°C for

2 min 30 s, followed by 5 min at 72°C after the last cycle. We then purified the PCR product using the Monarch PCR & DNA Cleanup Kit (NEB T1030), and sent it for Sanger sequencing (Microsynth AG, Switzerland) using primers pr078 and pr082 (Table S3).

*Curing the pKDsgRNA-attB vector.* We cured the thermosensitive vector pKDsgRNA-attB by re-growing candidate clones at 37°C without spectinomycin selection. We named one clone $MR^M$::dfrB9/pCas9-CR4 and used it in subsequent experiments.

**Creation of a variant library using gene editing**

*Preparing a parental strain.* We constructed a vector expressing guide RNA to target the chromosomal gene *folA*. We named this vector pKDsgRNA-folA. The construction principle is the same as for pKDsgRNA-attB, i.e., we replaced the gRNA spacer in pKDsgRNA-ack through circular polymerase extension cloning[44,99]. We performed two PCRs to amplify the vector backbone of pKDsgRNA-ack, using primer pairs pr009/pr049 and pr010/pr050 (Table S3). For these PCRs we used 0.020 U/µl Phusion DNA polymerase in Phusion HF buffer (NEB M0530) with 0.5 µM of each primer and 0.2 mM of each nucleotide. The thermocycling program consisted of 98°C for 3 min, 36 cycles of 98°C for 10 s, 55°C for 20 s, and 72°C for 1 min 30s, followed by 72°C for 5 min. Primers pr049 and pr050 contained overhangs with 20 nt sequences that are homologous to nucleotide positions 76-95 of *folA* (GenBank accession number NP_414590). We treated amplified fragments with 10 units DpnI and ExoI for one hour at 37°C, and purified them using the Monarch PCR & DNA Cleanup Kit. We combined 0.08 pM of each fragment using a single circular polymerase extension cloning reaction[99] that contained Phusion HF buffer, 0.02 U/µl of Phusion DNA polymerase, and 0.2 mM nucleotides. The reaction program consisted of 98°C for 30 s, 15 cycles of 98°C for 10 s, 55°C for 30 s, and 72°C for 2 min, followed by 72°C for 5 min. We added 2 µl of the reaction mixture to freshly prepared $MR^M$::*dfrB9*/pCas9-CR4 competent cells, which we subjected to electroporation (1 mm cuvette, 2.4 kV, BioRad MicroPulser) followed by 2 h recovery in SOC medium at 30 °C. We plated the transformed cells on SOC agar plates (50 µg/ml spectinomycin, 34 µg/ml chloramphenicol), and incubated them at 30°C for 40 h.

We verified the correct construction of pKDsgRNA-*folA* in three isolated colonies by Sanger sequencing (Microsynth AG, Switzerland) using the primer pBAD-rev (Table S3). In addition, we verified the activity of Cas9 using the Cas9-induced mortality assay. To this end, we resuspended bacterial colonies in PBS buffer, and serially diluted the suspension up to $10^5$-fold. We grew samples from these dilutions on SOB plates (34 µg/ml chloramphenicol and 50 µg/ml spectinomycin) with or without 0.16 µg/ml anhydrotetracycline. After 24 h incubation at 30°C we compared CFU counts on the anhydrotetracycline-induced and uninduced plates. We designated one confirmed clone as $MR^M$::*dfrB9*/pCas9-CR4/pKD-gRNA-*folA*, and used this clone later as a parental strain to create mutant libraries.

*Design of recombination template for gene editing*. We used a library of synthetic oligonucleotides as repair templates for gene editing[44,103]. To design optimal oligonucleotides, we used a custom script (https://gitlab.com/apson/templatelib). The script performs a search for candidate oligonucleotides and ranks the candidates according to parameters known to affect gene editing efficiency, such as GC content and secondary structure[81]. The script executes the following algorithm:

1. Select a target position in the genome to be mutated.

2. Create a collection of oligonucleotides that (i) have a sequence complementary to the genomic region in which the target lies, (ii) are 80-90 nts long, (iii) have a center that lies no farther than 10 nts from the target position.

3. Compute GC content, length, and secondary structure (ΔG) of oligonucleotides. (We calculated ΔG with the `seqfold` Python library[104].)

4. Score oligonucleotides by penalizing deviations from the following optimal parameter values*: GC content*=50%, *length*=80 bases, and *ΔG*=5 kcal/mole. For this scoring procedure, we increased the penalty score by (i) one for each one percent difference in *GC content*; (ii) three for each one kcal/mole decrease in *ΔG*; and (iii) one for each one nucleotide increase in *length*.

5. Rank oligonucleotides based on the sum of penalty scores. Select the top-ranked oligonucleotide, i.e., the oligonucleotide with the smallest penalty.

To design repair templates for many mutants, the oligonucleotides we designed in this way contained several degenerate positions (N), at which all four nucleotide bases A, C, G, T are incorporated with equal frequency during synthesis, generating all possible sequence combinations at that position. The degenerate oligonucleotide t022 used in this study is shown in Table S3. The oligonucleotide includes nine consecutive N bases corresponding to nucleotide positions 76-84 of the *folA* open reading frame (GenBank accession number NP_414590). These positions correspond to amino acid positions 26-28 of the DHFR protein. The total number of unique nucleotide sequences contained in the oligonucleotide library is $4^9$ (262,144), which corresponds to $21^3$ (9261) unique protein sequences (including stop codons). We also designed these oligonucleotides with three bases at the 5'-end and two bases at the 3'-end, which contain phosphorothioate bonds that provide protection from exonucleases[105]. We obtained the oligonucleotide library from IDT technologies (Belgium).

*CRISPR-Cas9 mutagenesis.* We prepared an overnight culture by inoculating a single colony of $MR^M$::*dfrB9*/pCas9-CR4/pKD-gRNA-*folA* into 1 ml SOB (34 µg/ml chloramphenicol, 50 µg/ml spectinomycin). After 24 hours of incubation at 30 °C and 225 rpm, we transferred 80 µl of the culture to 5 ml of fresh SOB medium (34 µg/ml chloramphenicol, 50 µg/ml spectinomycin), and incubated the culture at 30 °C with 225 rpm shaking. After five hours, we induced the lambda Red genes by adding 0.2% arabinose. Simultaneously, we induced the expression of *dfrB9* by adding 10 µM of vanillic acid. After 40 min, we harvested the induced culture to prepare electrocompetent cells using the glycerol-mannitol method. Next, we transformed the competent cells with 1mM of the oligonucleotide template t022 using a 1 mm cuvette (2.4 kV, BioRad MicroPulser). We allowed cells to recover in 1 ml of SOC medium at 30°C with shaking for two and a half hours, serially diluted the culture $10^1$-$10^3$-fold, and plated onto square plates (Thermo Scientific 240835; 245 mm x 245 mm). The plates contained SOC agar (34 µg/ml chloramphenicol, 50 µg/ml spectinomycin, 0.16 µg/ml anhydrotetracycline, 100 µM vanillic acid). After incubation at 30°C for 24 h, we scraped the bacterial lawn and resuspended it in PBS, centrifuged (3000 g, 20 min), and re-suspended again in 25 ml of PBS containing 15% of glycerol. We snap-froze the resuspended cells in liquid nitrogen and stored them at -80°C.

**Mass selection**

*Depleting DfrB9.* The cells stored at -80°C contain functional molecules of DfrB9, since we created the library by expressing *dfrB9* to ensure the viability of all mutants. The gene product of *dfrB9* can mask deleterious effects of *folA* mutations during a mass selection experiment. Therefore, we depleted the cellular pool of DfrB9 by growing the libraries without vanillic acid, a condition that represses *dfrB9* expression. We determined the time required for *dfrB9* depletion empirically by measuring the growth rate of depleted libraries after different depletion times (Fig. S22). Based on these results, we used a depletion time of 9 hours.

We thawed a library aliquot from -80°C on ice, and inoculated its cells into 40 ml of fresh M9 medium (0.4% glucose, 0.2% casamino acid) with no vanillic acid. The initial cell density was $OD_{600}$=0.088. We incubated the resulting culture at 30°C while shaking at 225 rpm for 9 h prior to mass selection.

*Selection.* After DfrB depletion, we performed a mass selection experiment in 25 ml of M9 medium containing 0.4 µg/ml trimethoprim using 250-ml polystyrene tissue flasks (Sarstedt 83.3911.002). We used 6 replicate cultures for each treatment, and incubated each replicate culture for 14 h at 30°C with shaking at 225 rpm.

We determined cell densities before and after selection by measuring absorbance ($\lambda$=600 nm, Fig. S23) in 1 cm cuvettes using a Genesys 180 UV-Vis spectrophotometer (ThermoFisher). In addition, we determined cell densities by diluting the cell cultures $10^4$, $10^5$, and $10^6$-fold and plating them on SOB agar with 10 µM of vanillic acid. After incubating at 30°C for 24 h, we counted the number of colony-forming units (CFUs). The average number of generations, as estimated with the formula $\log_2(\mathrm{CFU}_{\mathrm{after}}/\mathrm{CFU}_{\mathrm{before}})$, equaled 3.51 cell divisions.

*Amplicon sequencing.* We sampled cells from the mass selection experiment before and after selection to extract genomic DNA using a Dneasy Blood & Tissue Kit (Qiagen). Subsequently, we amplified the mutagenized region of *folA* from the genomic DNA using a combination of reverse primer ngsR1 (Table S3), and one of the six forward primers (ngsF01-ngsF06, Table S3). The forward primers contain 6 letter barcodes allowing us to pool amplification products. We performed amplification in 50 µl (0.02 U/µl Phusion DNA polymerase (NEB M0530), Phusion HF buffer, 0.2mM primers, and 0.2 nM nucleotides) with the following program: 98°C for 3 min, 36 cycles of 98°C for 10 s, 63°C for 20s, and 72°C for 30s, followed by 72°C for 5 min. After amplification, we treated PCR products with 10 U of DpnI and ExoI enzymes for 1 h at 37°C. We purified the PCR products using a Monarch PCR & DNA Cleanup Kit, pooled them equimolarly, and sent them for paired-end sequencing on the Illumina NovaSeq 6000 platform (Eurofins Genomics, Germany).

**NGS analysis pipeline and variant counting**

*Processing sequencing reads data.* We processed raw fastqc files using a custom analysis pipeline (gitlab.com/apson/ampseq_nextflow) written in `Nextflow`[106]. The pipeline includes the following steps:

1. Examine the quality of reads in fastqc files using `FastQ` with default parameters (v0.11.9, www.bioinformatics.babraham.ac.uk/projects/fastqc).
2. Demultiplex raw sequencing reads (using barcodes) and trim adapters with `cutadapt` version v3.4 (parameters “`-e 0.1 -no-indels -overlap=8`”)

[107]. In addition, `cutadapt` also filters out low quality reads. For this purpose, we used the parameter "`–max-ee=2`", which results in the removal of all trimmed reads with an expected error rate of more than two bases per read (see details in ref. [108]).
3. Merge paired reads using `flash` with the arguments "`-O -m 60 -M 140`" (v1.2.11)[109].
4. Align reads to the reference *folA* sequence using `bwa` (v0.7.17)[110] with high gap open penalties for deletions and insertions (`-O 16,16`). Sort and index aligned reads files using `samtools` (v1.7)[111].
5. Import aligned reads into the Python environment with the help of the python library `pysam` (v0.16.0) (github.com/pysam-developers/pysam). Filter out reads containing indels, call unique variants, and count variant frequencies.
6. Discard variants with fewer than 100 merged reads before selection. Export the frequency table of variants for subsequent analysis.

Table S4 shows pipeline statistics and read numbers for different samples.

We determined the threshold for variant detection empirically by comparing the number of variants in the mutagenized region, where most variants will be true variants, and outside the mutagenized region, where mutations are more likely to be sequencing errors. In other words, the number of mutations outside the mutagenized region provides a baseline level of false positive variants due to sequencing errors and possibly other experimental artifacts. We deliberately chose a conservative threshold for the inclusion of variants for further analysis. Specifically, we included only variants with more than 100 merged reads in all further analyses (Fig. S25), because our comparison of variant numbers inside and outside the mutagenized region indicated that variants above this threshold show fewer than 22 expected false variants per 261,382 detected variants (before selection), a false detection rate of 0.000085 (Fig. S25).

**Calculating fitness**

*Justification of the chosen method*

Different methods are used to express relative fitness in competition experiments[112]. One common method is to take the ratio of exponential growth rates for the variant of interest and the reference variant, $r_{variant}/r_{ref}$. In this case, the fitness value corresponding to the reference equals one. However, this ratio has the serious disadvantage that it does not directly relate to a selection coefficient[112,113]. Thus, it is preferable to use the difference $(r_{\text{variant}} - r_{\text{ref}})$, because according to well-established population genetic theory, it is this difference that corresponds to a selection coefficient[47]. In this representation, a variant with a growth rate equal to the reference has a relative fitness of 0. Because selection coefficients are important parameters for our analysis of adaptive evolution on a landscape, we implemented the second approach, as detailed in the next paragraphs.

*The model*

To estimate relative fitness, we used a well-established model of competition in a population of haploid genotypes (Chapter 1 in ref. [47]). The model assumes exponential growth, continuous time, overlapping generations, and no density- or frequency-dependent selection. Under these assumptions, the number $N$ of individuals of a given variant at time $t$ can be described as $N_t = N_0 e^{mt}$, where $N_0$ is the number of individuals at time 0, and $m$ is the variant-specific Malthusian parameter or intrinsic rate of increase. Fundamental population genetic theory shows that the relative frequency of variant $i$ competing with other variants changes linearly on the logit scale (logarithm of the odds ratio, see equations 1.6.8a and 1.6.12 on pages 26-27 in ref. [47]):

$$\ln\left(\frac{p_t}{1-p_t}\right) = \ln\left(\frac{p_0}{1-p_0}\right) + (r_i - \bar{r})t \quad (1)$$

where $p$ is the frequency of variant $i$ at the time indicated by the subscript, $r_i$ is the intrinsic rate of increase for variant $i$, and $\bar{r}$ is the average intrinsic rate of increase for all variants ($\bar{r} = r_1 \times p_1 + r_2 \times p_2 + \cdots + r_n \times p_n$). The expression $(r_i - \bar{r})$ is the relative rate of increase of variant $i$ during competition. It corresponds to the relative fitness[112]. If time $t$ is represented in units of generation time, we can convert $r$ to Wrightian fitness as $w = e^r$ (page 9 in [47]).

If we then denote the number of sequencings reads for variant $i$ as $N(i)$ and for all other variants as $N(other)$, we can replace the variant frequencies in equation 1 using the equality

$$\text{logit}(p) = \ln\left(\frac{p}{1-p}\right) = \ln\left(\frac{N(i)}{N(other)}\right)$$

to yield

$$\ln\left(\frac{N(i)_t}{N(other)_t}\right) = \ln\left(\frac{N(i)_0}{N(other)_0}\right) + (r_i - \bar{r})t \quad (2)$$

*Statistical estimation of relative fitness*

Based on equation 2, we can estimate the term $(r_i - \bar{r})$ by fitting a generalized linear model to sequencing read data using the logistic regression

$$Y = \beta_0 + \beta_1 t + \varepsilon,$$

where $Y$ is the dependent variable, which is given by the log odds ratio of sequencing read counts $\ln\left(\frac{N(i)}{N(other)}\right)$. The parameter $\beta_0$ is an intercept estimating $Y$ at time 0 (before selection), $\beta_1$ is a slope corresponding to $(r_i - \bar{r})$, $t$ is time, and $\varepsilon$ is an error term. We used a generalized linear model with the binomial family distribution because it uses the logit transformation (as a link function), which matches the logit-transformed sequencing read counts in equation 2.

To compare the fitness of any two mutants $i$ and $j$, we took advantage of the fact that $r_{ij} = r_i - r_j = (r_i - \bar{r}) - (r_j - \bar{r})$. Instead of fitting one model for each mutant, it is advantageous to compute the fitness difference $r_{ij}$ in a single step:

$$Y = (\beta_0 + \gamma * \beta_2) + (\beta_1 + \gamma * \beta_3)t + \varepsilon,$$

$$\gamma = \begin{cases} 1, \text{variant i} \\ 0, \text{variant j} \end{cases}$$

In this model, $\gamma$ is a dummy variable that equals one for variant *i* and zero for variant *j*. When $\gamma$ equals zero, we obtain the previous model with the intercept $\beta_0$ and slope $\beta_1$ for variant *j*. When $\gamma$ equals one, the coefficients $\beta_2$ and $\beta_3$ estimate the *difference* of the intercepts and slopes between the two variants. Therefore, the coefficient $\beta_3$ estimates the fitness $r_{ij}$ of variant *i* relative to variant *j*. The advantage of using this joint model is that it directly estimates the uncertainty for $r_{ij}$ (as a standard error for the $\beta_3$ term) and allows for a statistical test of the null hypothesis that $\beta_3 = 0$ (e.g., the Wald test).

We fitted the generalized linear models to the sequence read data for a given pair of variants with the built-in R function `glm`, using the argument "`family=binomial (link = logit)`." We obtained the estimates of model coefficients, standard errors, test statistics, and corresponding *p*-values using the built-in R function `summary.glm`.

With this procedure, we determined the fitness of each mutant in the library relative to the wild type variant. We adjusted *p*-values using the Benjamini-Hochberg false discovery rate method (261,331 tests). With the same procedure, we also calculated $r_{ij}$ for all pairs of variants that are one or two mutational steps apart (1 or 2 nucleotide substitutions), and tested the null hypothesis that $r_{ij} = 0$, i.e., implying that the fitness difference is not significant. We again adjusted the resulting *p*-values using the Benjamini-Hochberg method (45,730,826 tests).

**Assessment of reproducibility in mass selection experiments**

*Reproducibility between replicates within the same experiment.* To assess the reproducibility between replicates of the same experiment, we compared read count frequencies across replicates from the main selection experiment library (section *Mass selection*). We found that these frequencies were highly similar (minimum Pearson's correlation coefficient of fitness values between replicates: 0.9459 before selection and 0.9994 after selection, Fig. S1).

*Reproducibility across independent experiments.* To make sure that our method yields reproducible fitness estimates across experiments, we compared the relative fitness of multiple variants from the main selection experiment (section *Mass selection*) with that of the same variants from a smaller experiment. The libraries for the small experiment contained fewer DHFR variants (N=256), allowing to sequence them at higher coverage. We generated six small libraries (six gene editing procedures), as we explain next.

*Generation and mass selection of small libraries.* We performed gene editing mutagenesis using the protocol explained in the section *Creation of mutant library using gene editing*. We used the same parental strain $MR^M$::dfrB9/pCas9-CR4/pKD-gRNA-*folA* (section *Preparing a parental strain*). However, we used a different oligonucleotide template for recombination. This template (m023, Table S3) contained a reverse complement of the degenerated sequence GANNNN, encoding four codons at DHFR position 27 (aspartic or glutamic acid) and all possible 64 codons at position 28. The theoretical size of the library was 4×64=256 variants. Using this template, we independently transformed six cultures of the parental strain, producing six independent libraries. Next, we performed 6-fold mass selection experiment in M9 with 0.4 µg/ml trimethoprim, followed by genomic DNA extraction and amplicon sequencing as described for the main experiment (section *Mass selection*). Using our analysis pipeline (see sections *NGS analysis pipeline and variant counting* as well as *Calculating* *fitness*), we identified 256 variants and measured their fitness relative to the wild type. We compared the fitness values between this small experiment and the main experiment and found a high Pearson's $r$=0.972 (Fig. S2A).

Because we used only one large library in the main experiment (section *Creation of mutant library using gene editing*), we were concerned that spontaneous background mutations or mutations induced by the off-target activity of Cas9 may impact our fitness estimates. However, the comparison of fitness values obtained in the six independently produced libraries shows high consistency (the pairwise Pearson's correlation between the small libraries was $0.991<r<0.997$). This result demonstrates that our protocol for creating mutant libraries is reproducible across the repetitions of gene editing (Fig. S2B).

**The effect of folA-mix on the growth of low activity variants.**

To maintain low activity variants of DHFR, we integrated and expressed the gene *dfrB9* (see section *Chromosomal integration of dfrB9 using gene editing*). In addition, we evaluated an alternative strategy for maintaining low fitness variants, which does not involve *dfrB9.* Namely, we attempted to grow DHFR variants on folA-mix M9, which was designed to complement metabolic deficiencies of *folA* null mutants in previous studies[76,88–91].

*Creating control parental strain without dfrB9 integration (dfrB9-).* Instead of the parental strain with integrated *dfrB9*, we used the control strain $MR^M$/pCas9-CR4/pKD-gRNA-folA without this gene (Table S2). We obtained this control strain by transforming the original $MR^M$ strain with the pCas9-CR4 and pKD-gRNA-folA vectors (Table S1, Fig. S18).

*Generation of a library in the presence of folA-mix*. We performed gene editing using the parental strain $MR^M$/pCas9-CR4/ pKD-gRNA-folA (Table S2). We followed the same procedure as described in the section *Creation of a variant library using gene editing*, except that after gene editing, we plated the library onto SOB agar supplemented with folA-mix, so that variants lacking DHFR activity could grow.

To assess the potential of folA-mix medium to support the growth of low fitness DHFR variants, we randomly picked 48 colonies harboring library members, resuspended them in PBS, and inoculated them into 96-well plates (TPP 92096) containing 200 µl of either M9 or M9 folA-mix. We incubated the plates at 30 °C while shaking at 1000 rpm using a Stuart SI505 incubator (Cole-Parmer, UK). After 24 h, we measured the end-point optical density ($\lambda$=600 nm) using a Spark plate reader (Tecan). We compared the growth of each of the 48 library clones in the presence and absence of folA-mix. Consistent with previous studies, we found that folA-mix significantly improves the growth of low-fitness variants (Fig. S19A)[76,88–91].

*The effect of folA-mix on the uniformity of variant frequencies after gene editing.* We determined the distributions of variant frequencies in the library produced by gene editing in the folA-mix medium. To this end, we sequenced the library without further growth or selection. We performed sequencing and data analysis using the protocols described in the sections *Amplicon sequencing*, as well as *NGS analysis pipeline and variant counting*. We analyzed the frequency distribution of variants in this library and found that despite the fitness improvement provided by the folA-mix, the variant distribution in the library sample obtained using folA-mix was strongly skewed (Fig. S19, B and C). By comparison, the distribution of variants in the main library obtained after integrating and expressing *dfrB9* (without folA-mix) was more uniform. Thus, expressing *dfrB9* is superior for maintaining nonfunctional or low activity DHFR variants than using folA-mix (Fig. S19, B and C).

**The effect of dfrB9 integration on the reliability of fitness measurements.**

For the Cas9 mutagenesis of *folA*, we expressed *dfrB9* before commencing mutagenesis. Although this procedure provided a more uniform distribution of *folA* variants in the pre-selection library (Fig. S19C), we wanted to make sure that the expression of *dfrB9* does not interfere with subsequent fitness measurements. Additionally, the chromosomal integration of *dfrB9* into the parental strain may have had a pleiotropic effect on fitness by affecting neighboring genes. To determine whether *dfrB9* integration and expression causes substantial biases in fitness, we compared the fitness of variants in two small libraries, one with integrated *dfrB9* (*dfrB9+*) and the other without integrated *dfrB9* (*dfrB9-*). Apart from the *dfrB9* integration, the genomic background was identical for the two libraries.

*Generation of small libraries using dfrB9+ and dfrB9- parental strains*. As *dfrB9+* background, we used the same parental strain as for the main library ($MR^M$::dfrB9/pCas9-CR4/pKD-gRNA-*folA*, section *Preparing a parental strain*). As control *dfrB9-* strain, we used $MR^M$/pCas9-CR4/pKD-gRNA-*folA* (Table S2, section *Creating control parental strain without dfrB9 integration*). For each parental strain, we obtained a small library using gene editing (the protocol described in section *Creation of mutant library using gene editing*). We used the oligonucleotide m023 (Table S3), which encodes 256 variants (codon GAN at position 27 and codon NNN at position 28). Because this oligonucleotide template was restricted to four codons at the DHFR position 27 (aspartic or glutamic acid), it resulted in mostly functional variants of DHFR and allowed us to grow *dfrB9-* variants using the folA-mix SOB medium.

*Mass selection of dfrB9+ and dfrB9- small libraries*. We performed mass selection in M9 without folA-mix, followed by genomic DNA extraction, and amplicon sequencing as described for the main experiment (section *Mass selection*). We estimated the relative fitness of variants using our analysis pipeline (see sections *NGS analysis pipeline and variant counting*, as well as *Calculating fitness*). The relative fitness values were highly correlated between *dfrB9+* and *dfrB9-* cells (Pearson's correlation coefficient $r$=0.9367, Fig. S2C). Thus, by using tunable expression of the integrated *dfrB9*, we were able to improve library uniformity and maintain low fitness DHFR variants, while obtaining reproducible fitness estimates.

**The effect of the *mutL13* genotype on the reliability of fitness measurements**

In order to increase the efficiency of gene editing, we used a parental strain with impaired mismatch repair (*mutL13*)[80]. This strain grows more slowly than the wild type *mutL* strain (Fig. S24). We were thus concerned that the *mutL13* genotype can have a pleiotropic effect on fitness measurements in our main experiment. In addition, the *mutL13* genotype might bias fitness estimation by increasing the rate of background mutations. To address this concern, we created a small mutant library using the wild type version of the *mutL* gene, and compared the relative fitness of variants to the fitness found in the pilot experiment 1(*mutL13*).

*Creating control parental strain with the wild type mutL*. We prepared the parental strain $MR^S$::dfrB9 (Table S2), which harbors the wild type *mutL* gene but is otherwise isogenic to the parental strain used in the main experiment[80]. We integrated the *dfrB9* gene into this strain and transformed the strain with the vectors pCas9-CR4 and pKD-gRNA-folA required for gene editing of *folA*. To this end, we used the procedures described in the sections *Chromosomal integration of dfrB9 using gene editing* and *Preparing a parental strain*. However, instead of starting with the $MR^M$ *E. coli* strain (*mutL13*), we used the isogenic strain $MR^S$, which was created in a previous study by replacing the *mutL13* allele in the $MR^M$ genome with the wild type allele[80] (Fig. S18).

Generation of a small library using the control strain with the wild type mutL. We used the control parent ($MR^S$::dfrB9/pCas9-CR4/pKD-gRNA-*folA*) to prepare a small library of DHFR variants using the protocol described in the section *Creation of mutant library using gene editing*. We used oligonucleotide template m023, which encodes 256 variants (Table S3, the same template as used for the other small libraries).

*Mass selection of the control strain with the wild type mutL*. We used this library to perform mass selection in M9, and measured fitness as described in sections *Mass selection*, *NGS analysis pipeline and variant counting*, as well as *Calculating fitness*. The only difference in protocol was that we reduced the growth time in the mass selection experiment to 12 h, because the strain with the wild type *mutL* background grows faster. We performed this experiment in parallel to the experiments described in section *Mass selection of dfrB9+ and dfrB9-*. We compared the relative fitness of DHFR variants in the wild type *mutL* background with their fitness in the presence of the *mutL13* allele (the library *dfrB9+* from the parallel experiment). The fitness values were highly correlated (Pearson's $r$=0.9621, Spearman's ρ=0.9396, Fig. S2D), showing that the mutator allele does not strongly affect the fitness ranking of DHFR variants.

**Comparison of relative fitness from mass selection to the growth rate and resistance in single culture**

*Isolating individual variants.* We validated the fitness estimates from the main mass selection experiment with growth measurements obtained from *E.coli* populations expressing individual DHFR variants. To this end, we isolated 30 DHFR variants from the main library by randomly picking individual colonies. To cover a broad range of fitness values, we picked 15 colonies from the library before selection (enriched with low fitness variants) and 15 colonies from the library after selection (enriched with high fitness variants). After isolating these clones, we determined their *folA* sequence using Sanger sequencing. To this end, we extracted genomic DNA with the Dneasy Blood & Tissue Kits (Qiagen 69504), and amplified from this DNA the mutagenized *folA* gene region using a PCR with primers pr014 and pr015 (Table S3). We prepared the PCR reaction by mixing 0.020 U/µl Phusion DNA polymerase, Phusion HF buffer (NEB M0530), 0.5 µM primers, and 0.2 mM nucleotides. We used the following thermocycling protocol: 98°C for 3 min, 36 cycles of 98°C for 10 s, 55°C for 30 s, 72°C for 60 s, followed by 5 min at 72°C after the last cycle. We purified the PCR product using the Monarch PCR & DNA Cleanup Kit (NEB T1030), and sent the purified fragment for Sanger sequencing (Microsynth AG, Switzerland), using the primers pr014 and pr015 (Table S3).

*Measuring growth rate.* We determined the growth of individual variants in single cultures. To this end, we prepared an overnight culture of each variant in M9 containing 10 µM vanillate at 30°C with 225 rpm shaking for 20h. We diluted the overnight cultures 20-fold in

200 µl of M9 containing no vanillic acid and incubated them at 30°C with 225 rpm shaking for 9h to deplete DfrB9. After depletion, we adjusted the cell density to $OD_{600}$=0.05 and transferred 20 µl of cells to a 96-well plate containing 180 µl of M9 and 0.4 µg/ml of trimethoprim in each well. We incubated the plate for 16 h at 30°C and 1000 rpm in a Spark plate reader (Tecan), measuring the optical density (at $\lambda$=600 nm) at 5 min intervals. In this way, we measured the growth dynamics of each isolate in 4 replicated cultures that were split into 2 blocks. We used the resulting optical density data points to determine the maximal growth rate. To this end, we $\log_2$-transformed optical density values and scaled time in units of hours. We fitted multiple tangent lines to growth data of each culture using a sliding window of 20 data points and a step size of one (using the `lm` function built into R). The slope of the steepest line (maximum slope) is the maximal growth rate, which estimates the number of cell divisions per hour. These maximal growth rate values were highly correlated with the corresponding relative fitness estimates from the mass selection experiment (Fig. S2E, Pearson's $r$=0.993, $p$=2.3x$10^{-27}$).

_Resistance to trimethoprim._ We determined the resistance of 30 individual variants in single cultures. We prepared an overnight culture of each variant in M9 containing 10 µM vanillate at 30°C with 225 rpm shaking for 20h. We diluted the overnight cultures 20-fold in 200 µl of M9 containing no vanillate and incubated them at 30°C with 225 rpm shaking for 9h to deplete DfrB9. We inoculated depleted cultures using 1:1000 dilution into 96-well plates with 200 µl of M9 containing different trimethoprim concentrations (0.0625, 0.125, 0.25, 0.5, 1, 2,4, 8, 16, 32 µg/ml). We set up four replicate cultures at each trimethoprim concentration for each of the 30 variants. We incubated the plates at 30°C and 1000 rpm using a Stuart SI505 incubator (Cole-Parmer, UK). After 24 h, we measured the end-point optical densities ($\lambda$=600 nm) of the cultures using a plate reader (Spark, Tecan). To analyze resistance data, we plotted the end-point density values (a vertical axis) against the corresponding trimethoprim concentration (a horizontal axis), and obtained dose-response curves by applying linear interpolation (function `interp1`, from the package `pracma` in R). Then, using the function `trapz` from the same package, we performed numerical integration to determine the area under the dose-response curve (AUC). To normalize AUC values, we divided them by the largest value of AUC. The normalized AUC values are highly correlated with the corresponding relative fitness estimates from the mass selection experiment (Fig. S2F, Pearson's $r$=0.987, $p$=1.1x$10^{-23}$).

**Determining non-functional mutations**

Most variants had dramatically lower fitness than the wild type. They account for the most prominent bell-shaped part of the distribution of fitness effects (see Fig. 1B). We considered all such low fitness variants to express non-functional DHFR. To determine a fitness cut-off value for these non-functional variants, we fitted a mixture Gaussian model to the fitness values using the `mixEM` function from the `mixtools` R package[114], by setting the number of components (distributions) to `k=3`. The bell-shaped distribution of our data had an estimated mean of $\mu$=-0.721 and a standard deviation of $\sigma$=0.069. We then used this distribution's 0.999 quantile as a fitness cut-off value below which we considered a variant non-functional. This procedure resulted in a relative fitness cut-off of $(r_i - r_{\mathrm{WT}})$= -0.507774.

To validate this cut-off value, we compared it to the distribution of fitness effects in variants with nonsense mutations (Fig. S5). Specifically, we considered only variants with two stop-codon mutations (*N=752*) to minimize the chance of stop-codon read-through. These double nonsense variants had a mean fitness of $\mu$=-0.732, and 750 (99.73%) of the 752 variants had a fitness lower than the cutoff value. These numbers show that our cut-off value is conservative in excluding non-functional variants.

**Analysis of the fitness landscape**

Constructing a network of variants. In our analyses we assume that even the smallest fitness difference we measured are visible to selection, i.e., no effectively neutral mutations exist. This assumption is justified by the observation that in organisms with large effective population sizes like *E.coli* ($N_e$=$10^8$), even tiny fitness differences that would not appear significant in our experiment may be visible to natural selection[54].

Under this assumption, we built a network of variants (genotypes, nodes) with the help of the R `igraph` library[115] using the following algorithm:

1. Establish a list of neighboring variant pairs, which differ only at a single nucleotide (i.e., they have a genetic distance *d*=1)
2. Exclude pairs where both variants are non-functional (i.e. their fitness is below the cut-off)
3. Construct a network by connecting each pair of neighboring variants with a directed edge (arrow). Any one such edge corresponds to a fitness-increasing mutation, i.e., it points from the variant with lower fitness to the neighbor with higher fitness.
4. Extract the largest connected subgraph (or giant component[116]) of the network using the `components` function of `igraph` with the argument "`mode=weak`".

This connected subgraph of variants is the fitness landscape we analyze. It contains 135,178 variants and 324,044 edges between them. It includes all viable-viable and lethal-viable variant pairs. We allowed lethal-viable pairs, because starting from a variant with very low fitness, a beneficial mutation may create a viable variant. We did not allow edges between nonfunctional-nonfunctional pairs, because such mutations could not be traversed by an evolving population. All but eleven viable DHFR variants are a part of this landscape. Only 42% of all nonfunctional variants from the library are part of the landscape, but they still constituted most variants in the landscape (87%, 117159/135178).

Fitness peaks and accessible paths. We define a fitness peak as a variant (node) which has a higher fitness than all its neighbors.

Accessible paths. An evolutionary accessible path is a sequence of mutational steps (edges), in which every mutation increases fitness. We searched for the shortest accessible path from a given variant to a fitness peak using the `shortest_paths` function in `igraph`. Multiple alternative shortest paths might exist in a landscape. To perform an exhaustive search of all shortest paths, we used the function `all_shortest_paths`. If an accessible path exists between a given variant and a peak, we refer to this peak as an accessible peak.

Basins of attraction. The basin of attraction of a fitness peak comprises all variants for which a given peak is accessible. We refer to the basin's size as the number of variants in the basin. To determine the accessibility of fitness peaks from other variants in the landscape, we applied the function `distances` from `igraph` to our network representation of the landscape, in which each fitness-increasing mutation corresponds to a directed edge. This function computes the length of shortest paths from a set of vertices (such as all variants in the landscape) to another set of vertices (such as fitness peaks) . Each such length equals the number of mutational steps along the shortest accessible path from a variant to a fitness peak. The function also identifies vertex pairs with no accessible paths between them. The output of this function allowed us to determine the size of basins of attraction for each peak.

*Overlap between basins.* The basins of attraction of different fitness peaks may comprise overlapping sets of variants. To estimate the overlap between two basins $B_1$ and $B_2$, we used the Jaccard similarity coefficient $J$, which is equal to the size of the intersection between the two sets of variants divided by the size of their union:

$$J = \frac{B_1 \cap B_2}{B_1 \cup B_2}$$

*Detecting sign epistasis.* We detected pairwise epistatic interactions between mutants by enumerating all 4-variant motifs of variant pairs with genetic distance 2 (double mutants) in the landscape, using the `motifs` function from the `igraph` library in R. We classified each motif as belonging into one of the three categories: reciprocal sign epistasis, single sign epistasis, and no sign epistasis[11,117] (see Fig. S16A). The last category (no sign epistasis) includes both magnitude epistasis and additivity (no epistasis) without distinguishing them, because neither of the two subcategories affects peak accessibility[11,117].

*Extradimensional bypasses*

Reciprocal sign epistasis causes direct evolutionary paths between a low fitness variant *ab* and a higher-fitness double mutant *AB* to be non-accessible (Fig. S16C). However, a fitness landscape may contain indirect path(s) that lead from *ab* to *AB* and that are accessible. Such an indirect path is also called an extradimensional bypass[37,62,63,118]. For each case of reciprocal sign epistasis, we searched for extradimensional bypasses using the following algorithm:

1. Rank the variants in a four-variant motif by fitness, where rank 1 corresponds to the highest fitness. We arbitrarily denote the corresponding phenotype by *AB*. Because of reciprocal sign epistasis, its two-mutant neighbor *ab* will have the second-highest fitness (rank 2). The remaining variants *aB* and *Ab* will have rank 3 and 4 (see Fig. S16).
2. Search for the shortest accessible path, i.e., a path along which fitness increases in each step, from variant *ab* (rank 2) to variant *AB* (rank 1) within the whole network, using the `distance` function from the `igraph` library with the argument "`mode=out`". If such a path exists, this function computes the length of this path.

**Simulation of adaptive evolution on the landscape**

We assume that adaptive evolution on our landscape falls into the so-called strong selection weak mutation (SSWM) regime[55], because the probability that a mutation occurs in our focal nine nucleotide *folA* region in the *E. coli* genome is very low (9 positions × $2.2\times10^{-10}$ substitutions per position per generation)[56]. At this rate, any beneficial mutation is likely to become fixed before another mutant appears that will eventually go to fixation. In consequence, a population would usually harbor only one segregating mutation at a time. This means that adaptive evolution can be viewed as an adaptive walk, in which a population is monomorphic, and occupies a single genotypic location in the landscape, and then moves to an adjacent location via a single mutation-fixation event. From this new location, the next mutation-fixation event causes the population to move to another adjacent location, and so on, until the population reaches a fitness peak.

We initiated each adaptive walk by assigning a starting variant chosen at random from the landscape (see the next paragraph). We then calculated the fixation probability of each 1-mutant neighbor of this variant using Kimura's equation[119] (eq. 3.11 p.44): $F = (1 - e^{-2Nsp})/(1 - e^{-2Ns})$. In this equation, the fixation probability $F$ depends on the fitness effect $s$ of the focal neighbor (mutant), the initial frequency $p$ of the mutant, and the population size $N$. We used a population size similar to that of *E.coli*[54] ($N=10^8$) and assumed an initial frequency of $p=1/N$. After this calculation, we rescaled the fixation probabilities for all 1-mutation neighbors of the starting variant such that their sum equals one. We considered these rescaled probabilities as the values of a probability mass function for a multinomial distribution, which prescribes probabilistically which mutation reaches fixation next. For the first mutation-fixation event, we chose a 1-mutant neighbor of the starting genotype at random following this distribution. We then repeated this procedure for the new genotype, until the resulting adaptive walk reached one of the fitness peaks. To speed up simulations, we precomputed the multinomial distributions for each variant in the landscape.

We performed two kinds of simulations involving adaptive walks. In the first, we randomly selected $10^6$ starting variants by sampling them with uniform probability (and with replacement) from all variants in the landscape, excluding fitness peaks. For each of these $10^6$ starting variants we performed one adaptive walk. Because our landscape contains fewer than $10^6$ variants, we note that some variants would occur more than once as starting points of a random walk. We chose starting variants at random because we wanted them to be representative of all variants in the landscape in terms of properties such as fitness and adherence to a basin of attraction.

In the second case, we randomly selected 1000 variants from the landscape (without replacement) and performed 1000 simulations starting from each variant, which also yielded $10^6$ walks in total. Using a large number of adaptive walks that all start from the same variant is necessary to determine the extent to which the outcome of adaptive evolution is repeatable.

**Supplementary Note 1**

The expected number of peaks in a maximally rugged (uncorrelated) NK landscapes can be determined as the probability of a variant to be a peak times the number of variants in a landscape[13]. For our study this probability is equal to $1/[(A-1)\times L+1]=1/28$, where $A$=4 is the alphabet size and $L$=9 is the number of sites. The number of functional variants in our landscape is 18029. The expected number of peaks is thus $1/28\times 18029$, or 644 peaks.

**Supplementary Note 2**

Reciprocal sign epistasis creates local fitness reductions along mutational paths[11], it can render such paths inaccessible for Darwinian evolution. However, in a high-dimensional landscape, such local fitness reductions can be circumnavigated via extradimensional bypasses[37,62,63,79,118]. These are indirect evolutionary paths that require additional mutational steps to detour around a local fitness decrease, and are thus longer than direct (but inaccessible) paths (Fig. S16C). In our landscape, such paths do exist, but they can overcome only 23% of local fitness reductions caused by reciprocal sign epistasis (Fig. S16D). More importantly, 83.6% of shortest accessible paths from a DHFR variant to a high fitness peak are direct paths, i.e., paths as short as the genetic distance from the variant to the peak. These observations show that an abundance of extradimensional bypasses around a local fitness reduction is not necessary to explain the high navigability of our landscape.

**Supplementary References**


80. Sprouffske, K., Aguilar-Rodríguez, J., Sniegowski, P. & Wagner, A. High mutation rates limit evolutionary adaptation in Escherichia coli. *PLOS Genet.* **14**, e1007324 (2018).
81. Reisch, C. R. & Prather, K. L. J. Scarless Cas9 Assisted Recombineering (no-SCAR) in Escherichia coli, an Easy-to-Use System for Genome Editing. *Curr. Protoc. Mol. Biol.* **117**, 31.8.1-31.8.20 (2017).
82. Meyer, A. J., Segall-Shapiro, T. H., Glassey, E., Zhang, J. & Voigt, C. A. Escherichia coli “Marionette” strains with 12 highly optimized small-molecule sensors. *Nat. Chem. Biol.* **15**, 196–204 (2019).
83. Gentile, C. F., Yu, S.-C., Serrano, S. A., Gerrish, P. J. & Sniegowski, P. D. Competition between high- and higher-mutating strains of Escherichia coli. *Biol. Lett.* **7**, 422–424 (2011).
84. Liberfarb, R. M. & Bryson, V. Isolation, characterization, and genetic analysis of mutator genes in Escherichia coli B and K-12. *J. Bacteriol.* **104**, 363–375 (1970).
85. Warren, D. J. Preparation of highly efficient electrocompetent Escherichia coli using glycerol/mannitol density step centrifugation. *Anal. Biochem.* **413**, 206–207 (2011).
86. Zhu, L. J., Holmes, B. R., Aronin, N. & Brodsky, M. H. CRISPRseek: A Bioconductor Package to Identify Target-Specific Guide RNAs for CRISPR-Cas9 Genome-Editing Systems. *PLoS ONE* **9**, (2014).
87. Guo, J. *et al.* Improved sgRNA design in bacteria via genome-wide activity profiling. *Nucleic Acids Res.* **46**, 7052–7069 (2018).
88. Herrington, M. B. & Chirwa, N. T. Growth properties of a folA null mutant of Escherichia coli K12. *Can. J. Microbiol.* **45**, 191–200 (1999).
89. Krishnan, B. R. & Berg, D. E. Viability of folA-null derivatives of wild-type (thyA+) Escherichia coli K-12. *J. Bacteriol.* **175**, 909–911 (1993).
90. Hamm-Alvarez, S. F., Sancar, A. & Rajagopalan, K. V. The presence and distribution of reduced folates in Escherichia coli dihydrofolate reductase mutants. *J. Biol. Chem.* **265**, 9850–9856 (1990).
91. Singer, S., Ferone, R., Walton, L. & Elwell, L. Isolation of a dihydrofolate reductase-deficient mutant of Escherichia coli. *J. Bacteriol.* **164**, 470–472 (1985).
92. Howell, E. E. Searching Sequence Space: Two Different Approaches to Dihydrofolate Reductase Catalysis. *ChemBioChem* **6**, 590–600 (2005).

93. Faltyn, M., Alcock, B. & McArthur, A. Evolution and Nomenclature of the Trimethoprim Resistant Dihydrofolate (dfr) Reductases. (2019) doi:10.20944/preprints201905.0137.v1.
94. Gibson, D. G. *et al.* Enzymatic assembly of DNA molecules up to several hundred kilobases. *Nat. Methods* **6**, 343–345 (2009).
95. Kunjapur, A. M. & Prather, K. L. J. Development of a Vanillate Biosensor for the Vanillin Biosynthesis Pathway in E. coli. *ACS Synth. Biol.* **8**, 1958–1967 (2019).
96. Campbell, A. M. Chromosomal insertion sites for phages and plasmids. *J. Bacteriol.* **174**, 7495–7499 (1992).
97. Diederich, L., Rasmussen, L. J. & Messer, W. New cloning vectors for integration into the λ attachment site attB of the Escherichia coli chromosome. *Plasmid* **28**, 14–24 (1992).
98. Balbás, P. & Gosset, G. Chromosomal editing in Escherichia coli. *Mol. Biotechnol.* **19**, 1–12 (2001).
99. Quan, J. & Tian, J. Circular Polymerase Extension Cloning of Complex Gene Libraries and Pathways. *PLOS ONE* **4**, e6441 (2009).
100. Jiang, W., Bikard, D., Cox, D., Zhang, F. & Marraffini, L. A. RNA-guided editing of bacterial genomes using CRISPR-Cas systems. *Nat. Biotechnol.* **31**, 233–239 (2013).
101. Doench, J. G. *et al.* Rational design of highly active sgRNAs for CRISPR-Cas9-mediated gene inactivation. *Nat. Biotechnol.* **32**, 1262–1267 (2014).
102. Li, Y. *et al.* Metabolic engineering of Escherichia coli using CRISPR–Cas9 meditated genome editing. *Metab. Eng.* **31**, 13–21 (2015).
103. Jiang, Y. *et al.* Multigene Editing in the Escherichia coli Genome via the CRISPR-Cas9 System. *Appl. Environ. Microbiol.* **81**, 2506–2514 (2015).
104. Timmons, J. & leshane. Lattice-Automation/seqfold 0.7.7. (2021) doi:10.5281/zenodo.4579886.
105. Nikiforov, T. T., Rendle, R. B., Kotewicz, M. L. & Rogers, Y. H. The use of phosphorothioate primers and exonuclease hydrolysis for the preparation of single-stranded PCR products and their detection by solid-phase hybridization. *PCR Methods Appl.* **3**, 285–291 (1994).
106. Di Tommaso, P. *et al.* Nextflow enables reproducible computational workflows. *Nat. Biotechnol.* **35**, 316–319 (2017).
107. Martin, M. Cutadapt removes adapter sequences from high-throughput sequencing reads. *EMBnet.journal* **17**, 10–12 (2011).
108. Edgar, R. C. & Flyvbjerg, H. Error filtering, pair assembly and error correction for next-generation sequencing reads. *Bioinformatics* **31**, 3476–3482 (2015).

109. Magoč, T. & Salzberg, S. L. FLASH: fast length adjustment of short reads to improve genome assemblies. *Bioinformatics* **27**, 2957–2963 (2011).
110. Li, H. & Durbin, R. Fast and accurate short read alignment with Burrows-Wheeler transform. *Bioinforma. Oxf. Engl.* **25**, 1754–1760 (2009).
111. Danecek, P. *et al.* Twelve years of SAMtools and BCFtools. *GigaScience* **10**, giab008 (2021).
112. Chevin, L.-M. On measuring selection in experimental evolution. *Biol. Lett.* **7**, 210–213 (2011).
113. Blanquart, F. & Bataillon, T. Epistasis and the Structure of Fitness Landscapes: Are Experimental Fitness Landscapes Compatible with Fisher's Geometric Model? *Genetics* **203**, 847–862 (2016).
114. Benaglia, T., Chauveau, D., Hunter, D. R. & Young, D. S. mixtools: An R Package for Analyzing Mixture Models. *J. Stat. Softw.* **32**, 1–29 (2010).
115. Csardi, G. & Nepusz, T. The igraph software package for complex network research. *InterJournal* **Complex Systems**, 1695 (2006).
116. John, C. & Allan, H. D. *A First Look at Graph Theory*. (Allied Publishers, 1995).
117. Greene, D. & Crona, K. The Changing Geometry of a Fitness Landscape Along an Adaptive Walk. *PLOS Comput. Biol.* **10**, e1003520 (2014).
118. Cariani, P. A. Extradimensional bypass. *Biosystems* **64**, 47–53 (2002).
119. Kimura, M. *The neutral theory of molecular evolution*. (Cambridge University Press, 1983).
